# High avidity phase-separated RNA-protein sialogranules sense lectins and inhibit influenza infection

**DOI:** 10.64898/2026.02.22.707264

**Authors:** Or Willinger, Naor Granik, Tai Salomon, Sarah Goldberg, Roee Amit

## Abstract

We previously developed synthetic RNA-protein (sRNP) granules as stable, programmable, genetically encoded nanoparticles. Here, we adapt this platform into high-avidity sialogranules that inhibit influenza virus entry. Natural or synthetic α2,6-sialylated proteins and peptides were fused to RNA-binding domains and assembled with synthetic long non-coding RNA to form phase-separated glyco-nanoparticles. Using the sialic-acid-binding lectin Sambucus nigra agglutinin (SNA), we identified a selective transition from sialogranules to tri-component multiphasic biocondensates that occurs only for sialylated constructs. The structural features of this transition quantitatively correlate with predicted sialylation density, enabling extraction of an effective avidity constant. Enzymatic desialylation with neuraminidase or Endo H abolished SNA binding and restored native granule morphology. Guided by this assay, LAMP1-based sialogranules were selected and inhibited influenza entry by ∼50% in cell culture. These results establish sRNP sialogranules as programmable glyco-nanoparticles integrating glycan sensing with antiviral decoy activity.

## Introduction

Weak specific interactions are central to many biological processes relevant to health and disease^1^. A prominent example is lectin-glycan binding, which is intrinsically low affinity but achieves functional specificity through multivalent presentation of both lectins and glycans^1–3^. Such interactions underlie diverse processes including cell-cell communication, signal transduction, immune modulation during viral infection, bacterial pathogenicity, and viral entry^1,4–7^. Pandemic-potential human influenza viruses (IFV) exemplify this principle: hemagglutinin (HA) binds α2,6-linked sialic acid (SA) on host glycoproteins with weak affinity (Kd ∼ mM)^8^. To facilitate cell entry, the virion utilizes several hundred^9^ HA proteins to create a multivalent bond, generating a high-avidity interaction with sialylated receptors^2^. This multivalent avidity dramatically reduces virion dissociation rates from the cell surface, thereby enabling productive infection.

Given the ability of multivalency to increase the effective binding strength between low-affinity ligand and target^10^, engineering high avidity glyco-nanoparticles (glycoNPs) for diagnostical, translational, and therapeutic purposes has become a vibrant area of research^3^. Broadly, glycoNPs are typically composed of either gold-nanoparticles (GNPs), superparamagnetic iron oxide nanoparticles (SPIONs), polymeric nanoparticles (e.g. polyethyleneglycol (PEG) functionalized systems), or hybrid glycoNPs which combine a polymer with an inorganic core^3,11–13^. Recent examples include developing glyco-dendri-protein-nano-particles that are capable of blocking infection of T-lymphocytes and human dendritic cells by a model virus displaying Ebola glycoproteins^14^, and gold nano-particles (GNP) conjugated to glycans that inhibited DC-SIGN-mediated augmentation of Ebola virus glycoprotein-driven cell entry^15^. However, glycoNPs engineering faces several challenges that are currently limiting progress. First, for *in vitro* applications there is a need to maintain uniform glycan density, multivalency, orientation, and accessibility on NPs, which are necessary for achieving consistent biochemical performance^3,16–19^. Second, for *in vivo* applications, nanoparticles must evade rapid clearance, exhibit colloidal stability in biological media or sera, and enable sufficient glycan complexity to overcome biological heterogeneity in many target systems (e.g. tumors)^19^.

One possible approach to overcoming these challenges is to engineer synthetic biomolecular phase-separated biocondensates that can display a desired ligand or target in high density and functional state, creating multivalent biologically compatible and structurally flexible nanoparticles. We recently showed that we can create synthetic gel-like phase-separated biocondensates termed synthetic RNA-protein (sRNP) granules both in vitro and in cellulo in various cell types using programable synthetic long non-coding RNA (slncRNA) molecules featuring a cassette of connected hairpin structures, and a matching RNA binding protein (a tandem-dimer PP7-coat protein fused to mCherry, abbreviated as tdPCP-mCherry)^20,21^. We further showed in a separate study that these granules can be functionalized to become anti-SARS-CoV-2 multivalent decoy nanoparticles by fusing the soluble form of human ACE2 (hACE2) to the tdPCP moiety^22^. The decoy granules stored the soluble hACE2 receptor in high density; demonstrated long-term colloidal stability in both ambient room-temperature conditions and within biological media or animal’s blood-serum; sponge-like functionality when mixed with the SARS-CoV-2 receptor binding domain moiety^22^ or coronavirus-mimicking polystyrene beads^23^; and, a strong efficacy in inhibiting infection of both the Delta (B.1.617.2) and Omicron (BA.1) SARS-CoV-2 variants using a plaque entry assay^22^.

In this work, we adapt our synthetic RNP granule platform to form multivalent sialylated nanoparticles called sialogranules. To do so, we fused the tdPCP-mCherry backbone to naturally sialylated proteins that have either N-linked or O-linked glycosylation sites, and have been used previously as in-vitro influenza targets^24–28^. In addition, we also fused the tdPCP-mCherry backbone to unstructured synthetic peptides that were encoded with several instances of the N-X-(S/T) motif needed for N-linked glycosylation^29^. We then used the lectin *Sambucus nigra* agglutinin (SNA) to induce morphological changes in the candidate sialogranules as a function of avidity. This allowed us to differentiate between different sialogranules based on the sialic acid content of their protein component. Finally, we showed that the highest avidity sialogranule detected by our assay (LAMP1) inhibited IFV infection in an entry assay. Consequently, we show that the phase-separated RNA-protein sialogranules form high-avidity Velcro-like glycoNPs that can be used as both a sensitive lectin sensor and as a potential prophylactic that can inhibit glycan-mediated infections.

## Results

### Synthetic recombinant sialoprotein candidates form sialogranules in the presence of slncRNA cassettes

Building on the success of using the granule platform to screen for anti-SARS-CoV-2 functionality^22^, we sought do the same for influenza virus (IFV). Unlike the SARS-CoV-2 virus, which binds a single, well-defined receptor (ACE2), IFV targets post-translational modifications which are present on various host proteins and receptors. In particular, human IFV targets sialic acid (SA) attached to host glycoproteins via an α2,6 glycosidic bond^8^. SA is added to proteins in mammalian cells in various O-linked glycosylation (OLG) patterns and can also be found in two out of the three major N-linked glycosylation (NLG) patterns, namely the hybrid and complex structures^29^. To screen for anti-IFV functionality, we hypothesized that our phase-separated gel-like granules containing slncRNA molecules and a high density of RNA-binding sialylated proteins (sialogranules) can potentially form a high avidity target, and thus be used to detect binding of SA by a cognate lectin in a highly sensitive manner. For the third molecule capable of interacting with the SA modifications (mimicking the IFV hemagglutinin), we opted to use commercially-available FITC-labeled *Sambucus nigra* agglutinin (SNA), which has a stronger affinity to α2,6 SA than the IFV hemagglutinin protein^26,30,31^. Upon introducing SNA, we expected to observe either co-localization with the granules, or transition from the granule state to another phase-separated state involving all three components due to the known agglutination properties of SNA^32,33^.

To design candidate high-avidity anti-IFV sialogranules, we first identified a small set of putative properly sialylated proteins (i.e., possess α2,6 SA). The shared RNA-binding moiety of the granules’ different protein components, tdPCP, itself contains two occurrences of the amino acids N-S-T, meeting the requirements for the known NLG motif N-X-S/T (N=asparagine, X≠proline, S=serine, T=threonine). As a result, a granule composed from the slncRNA together with a tdPCP-mCherry moiety produced in mammalian cells (Figure 1A, left) provides a candidate sialogranule, while granules with the same slncRNA and with unsialylated tdPCP-mCherry produced in bacterial cells^34^ can serve as a negative control. Given the small number of theorized sialylated sites on tdPCP-mCherry, we opted to construct an additional variant, in which the tdPCP-mCherry protein is augmented by a synthetic peptide containing five N-X-S/T sites (NXST1-mCherry-tdPCP) (Figure 1A, right). We then expressed and purified the synthetic sialoprotein variants in both mammalian and bacterial cells (HEK293F and *E. coli* TOP10, respectively, see Methods), yielding a total of four slncRNA-binding variants: tdPCPb, tdPCPm, NXST1b, NXST1m.

**Figure 1.**
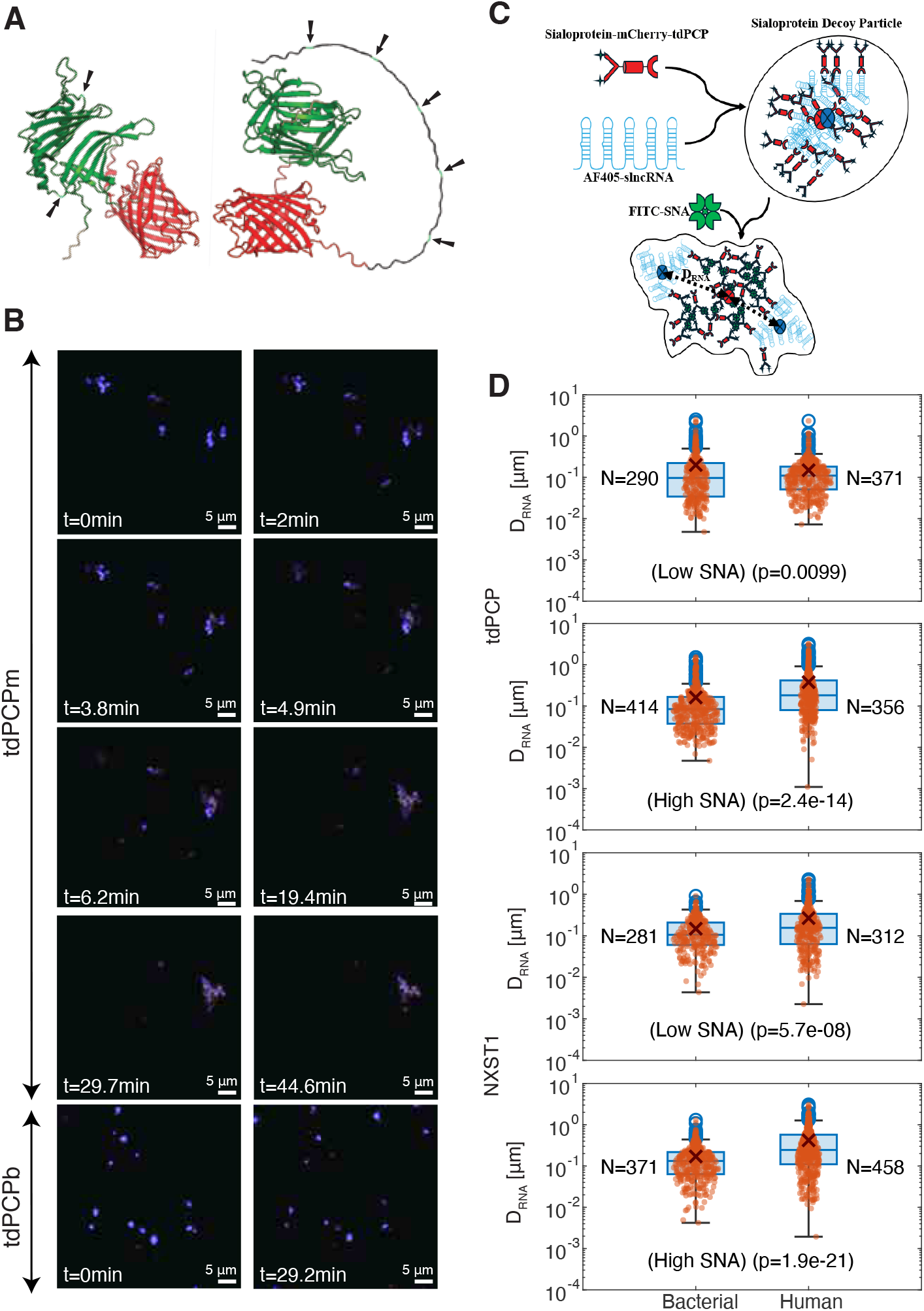
Phase separation-based agglutination assay identifies sialylated proteins. **(A)** AlphaFold structures for tdPCP (left) and NXST1 (right). Black, red, and green are NXST1, mCherry, and tdPCP domains, respectively. Putative sialylation sites are marked by arrows and light blue shading (asparagine residues in an N-X-S/T motif). **(B)** Accumulation of SNA signal (green) over time in a candidate sialogranules (red-blue). (Top) Candidate sialogranules composed of slncRNA (blue) and tdPCPm (red) transition to a multiphasic biocondensate composed of SNA, tdPCPm, and slncRNA after 20min incubation with SNA. (Bottom) granules composed of slncRNA and tdPCPb (red) do not transition to a multiphasic biocondensate. Scalebar: 5µm. Images are taken from Supplementary Movies S2 and S1, respectively. **(C)** Schematic model for the transition from a sialogranules (top) to a three-component multiphasic agglutinated biocondensate, where the sialoprotein and SNA form a dense liquid-like phase in the center, while the slncRNA segregates to the periphery of the structure over a distance *D*_*RNA*_. Circles mark the center-of-gravity of sialoprotein (red) and slncRNA (blue). **(D)** Distributions comparing *D*_*RNA*_ as defined in panel (C) for granules composed of bacterial-(left) and mammalian-(right) expressed proteins for low and high SNA concentrations. tdPCP – top two panels. NXST1 – bottom two panels. The mean of each distribution is marked by dark red X. Each red dot represents one mCherry-positive event for which the *D*_*RNA*_ was calculated. The student’s t-test was applied to ascertain statistical significance. Low and high SNA concentrations were set at below 1µg/ml (molar ratio of ∼1:50 SNA:sialoproteins) and above 500µg/ml (molar ratio of ∼3:2 SNA:sialoproteins), respectively.

To test for detection of sialylation, immediately after forming the granules we added the FITC-labelled SNA and observed the granules over 30min (Figure 1B). For granules containing the bacterially produced protein variants tdPCPb and NXST1b, no changes were observed during the 30min span of the experiment (Supplementary Movie S1). However, putative sialogranules containing tdPCPm and NXST1m exhibited an abrupt transition from a spot-like granule form to a lower-density, cloudy biocondensate containing all three components, within five minutes (Figure 1B, Supplementary Movie S2). A close examination of the images and movies reveal that this biocondensate is characterized by co-localization of SNA and sialylated protein in the center of the new biocondensates, while the slncRNA is found at the periphery of the cloud (Figure 1C). In addition, the movies and images also reveal that the new biocondensate appears to be assembled in a manner reminiscent of a nucleation process. Importantly, slncRNA and SNA did not colocalize without the presence of a sialoprotein (Figure S1A), and SNA-sialoprotein biocondensates were better detected when slncRNA cassettes were present in the reaction (Figure S1B).

Next, given the segregation of the slncRNA to the periphery of the new multiphasic SNA-agglutinated biocondensates, we posited that we could harness the slncRNA exclusion process to extract a quantitative observable for differentiation between SNA-agglutinated and non-SNA-agglutinated biocondensates. For this purpose, we define *D*_*RNA*_ as the distance between the center of slncRNA clusters and putative sialoprotein clusters (Figure 1C). We then extracted *D*_*RNA*_ from several hundreds of microscopic biocondensate events for each granule variant, in both low and high SNA concentrations (Figure 1D). For both the tdPCP and the NXST1 proteins, a statistically significant difference in *D*_*RNA*_ distributions was observed between granules containing bacterial and human-cell expressed proteins. Specifically, for low SNA concentrations the difference in mean *D*_*RNA*_ was found to be small and marginally significant for both tdPCPm (t-test p-value 1e-2) and NXST1 (t-test p-value 5.7e-8). However, for high SNA concentrations the average *D*_*RNA*_ observed for both the tdPCPm and NXST1m was found to be significantly larger as compared with tdPCPb (p-value=2.4e-14, student t-test) and NXST1b (p-value=1.9e-21, student t-test), respectively. The increase in average *D*_*RNA*_ is consistent with a transition from a granular structure to a new type of multiphasic biocondensate, characterized by a central region that is dominated by a dense matrix of agglutinated SNA and sialoprotein and condensed slncRNA that is segregated to the periphery (see Figure 1C for structural model). Consequently, detection of sialic acid on granule binding protein is possible via characterization of a structural transition from the granular phase to a multiphasic agglutinated biocondensate (MAB) phase upon introduction of SNA.

### Recombinant sialoprotein candidates form functional sialogranules in the presence of slncRNA cassettes

We next considered four natural human proteins that were either known to be sialylated, or were used previously in influenza research^24–28^: LAMP1 (Figure 2A), GYPA (Figure S2A), fetuin (Figure S2B) and ACE2. After fusion of the soluble domain to the tdPCP moiety, these candidate proteins contained a putative set of 21, 15, 4 and 9 sialic acid modification sites, respectively. Since protein glycosylation is sometimes crucial for the proper folding of a protein^35^, no bacterial versions of these proteins were made. Additionally, we created two additional synthetic sialoprotein candidates, comprised of tandem repeats of the NLG sequon: NXST2m, similar to the previously described NXST1m but with different amino acids in the X position (see Figure S3D), and tdNXST1m, a tandem repeat of NXST1m (Figure S3D). Once fused to tdPCP, these two additional synthetic sialylated protein candidates contain 7 and 12 theoretical sialylated sites, respectively. All of these additional candidates formed granules when incubated with the slncRNA, and formed MABs in the presence of SNA that were similar to the biocondensates observed for tdPCPm and NXST1m (i.e. where the RNA was found to cluster at the periphery, Figure 1), indicating that a similar phase separation mechanism is likely acting on all systems (Figure 2B, Figure S2E, and Supplementary Movie S3).

**Figure 2.**
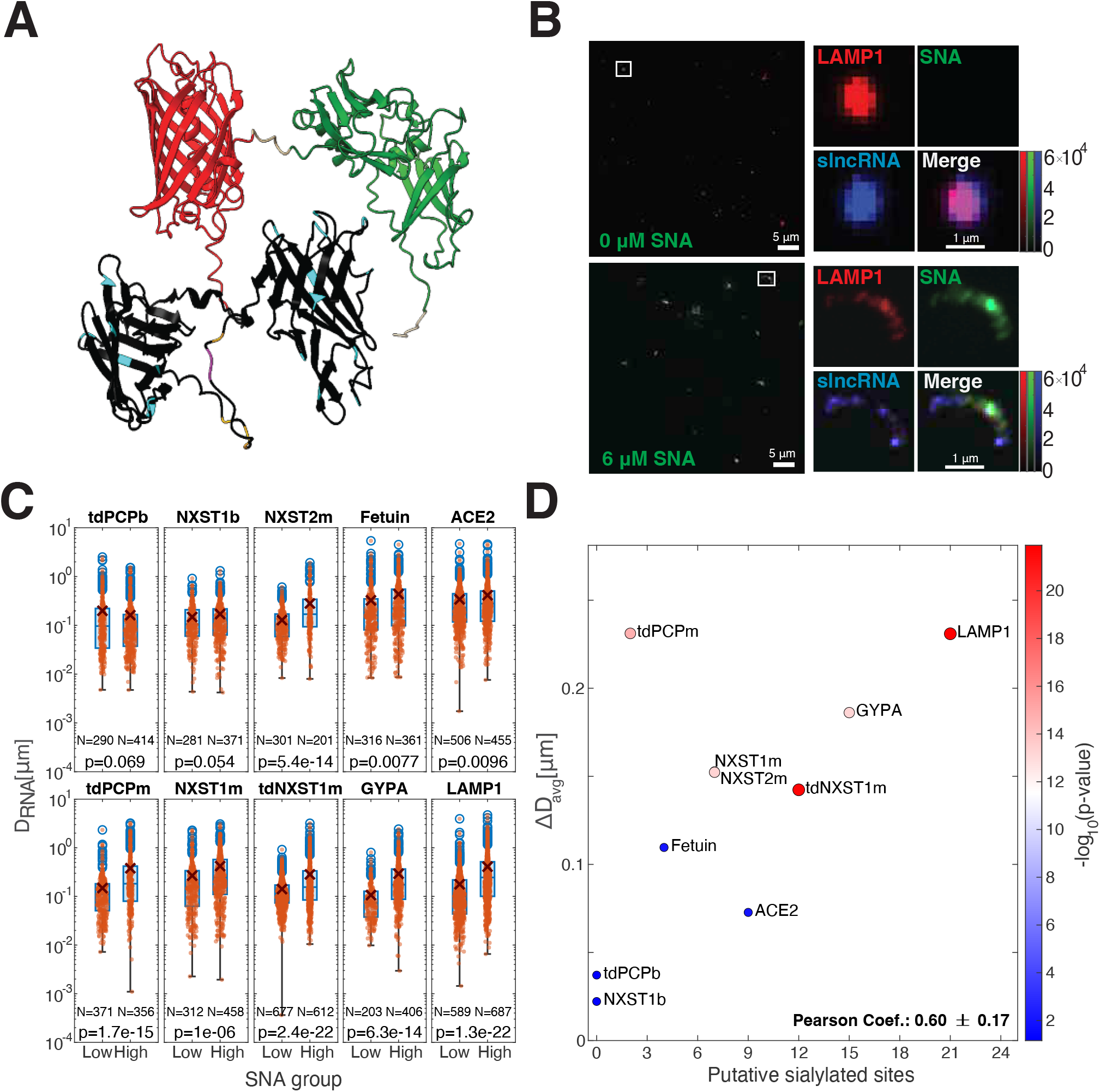
SNA-dependent formation of multiphasic agglutinated biocondensates (MABs). **(A)** Predicted AlphaFold structure for LAMP1-mCherry-tdPCP. Black, red, and green are LAMP1, mCherry, and tdPCP domains, respectively. Putative sialylation sites are marked by light-blue shading (asparagine residues in an N-X-S/T motif), yellow (serine residue), and magenta (threonine residue). **(B)** Typical field-of-view of LAMP1 granule (top) and the MABs (bottom) divided into individual channels. (Top-left) protein (mCherry). (Top-right) SNA (AF488). (Bottom-left) slncRNA (AF405). (Bottom-right) Merged panel with all three images overlayed. (Top-set) No SNA added (scalebar: 5µm FoV, 1µm enlargement). (Bottom-set) High SNA concentration, corresponding to the 2mg/ml stock (scalebar: 5µm). **(C)** Box-plot distributions for *D*_*RNA*_ for each candidate sialoprotein. The mean of each distribution is marked by dark red X. Each red dot represents one single mCherry-positive event for which the *D*_*RNA*_ was calculated. Low and high SNA concentrations were set at below 1µg/ml (molar ratio of ∼1:50 SNA:sialoproteins) and above 50µg/ml (molar ratio of ∼3:2 SNA:sialoproteins), respectively. The student’s t-test was applied to ascertain statistical significance. **(D)** Δ*D* as a function of putative sialylated sited. Δ*D* is defined as the difference between the mean *D*_*RNA*_ values for high and low SNA concentrations obtained for each protein in panel C. Color and size of circle correspond to the student’s t-test p-value computed for each panel in Figure 2C. We note that NXST1m and NXST2m appear to have nearly the same Δ*D* value and have the same number of putative sialylated sites.

We then quantified the *D*_*RNA*_ values for several hundred biocondensates at low and high SNA concentrations, for all of the additional candidate sialylated proteins (Figure 2C). The results show large statistically significant differences in the *D*_*RNA*_ distributions between condensates observed in the presence of low and high SNA concentrations for LAMP1 (p-value=1.3e-22 student, t-test), GYPA (p-value=6.3e-14, student t-test), tdNXST1m (p-value=2.4e-22, student t-test), NXST2m (p-value 5.4e-14, student t-test), and tdPCPm (p-value=1.7e-15, student t-test). Three proteins exhibited a moderate statistically significant difference in *D*_*RNA*_ distributions between the low and high SNA states with a low degree of statistical significance: NXST1m (p-value=1e-6, student t-test), fetuin (p-value=7.7e-3, student t-test) and ACE2 (p-value=9.6e-3, student t-test). Finally, the bacterial proteins (tdPCPb and NXST1b) were found to have no statistically significant difference in *D*_*RNA*_ distributions (p-value=0.069 and p-value=0.054, respectively, student t-test) between the high and low SNA states, as expected.

Finally, we plotted the difference between the distribution means for low and high concentrations of SNA (Δ*D*_*RNA*_) for each protein in our set as a function of number of putative sialylated sites (Figure 2D) and found them to be strongly correlated (Pearson coefficient = 0.6). Interestingly, tdPCPm, exhibiting a much larger Δ*D*_*RNA*_ response as compared with the general trend, suggesting that the number of sialylated sites or sites that are available for interaction may be influenced by other features^36^. In addition, both tdPCP and NXST1 exhibited a ∼6-7-fold difference in Δ*D*_*RNA*_ between the mammalian and bacterial isoforms (tdPCP: bacterial=37.1nm, mammalian=231nm, NXST1: bacterial=22.1nm, mammalian=152.9nm), providing further evidence that the underlying response might be dependent on sialylation.

### Kinetics of both SNA binding to sialogranules and slncRNA exclusion exhibit a dependence on the number of putative sialylation sites

Given the correlation observed between the number of putative sialylated sites and Δ*D*_*RNA*_, we wondered whether varying the SNA concentration could yield a typical dose-response-like kinetic transition from the sialogranule state to the MAB state. To check this, we mixed a pre-formed fixed amount of granules with a set of ten increasing SNA concentrations, and analyzed the phase-separated structures that formed. In brief (see Methods), for each of the sialoprotein candidates (see Figure 3A for a sample field of view), we identified mCherry-positive events that were colocalized with either SNA or slncRNA (Figure 3B) and classified them into two analysis groups based on their intensity and percent of overlap of the mCherry signal and either FITC or AF405 signals, respectively (Figure 3C, diagram, see Methods).

**Figure 3.**
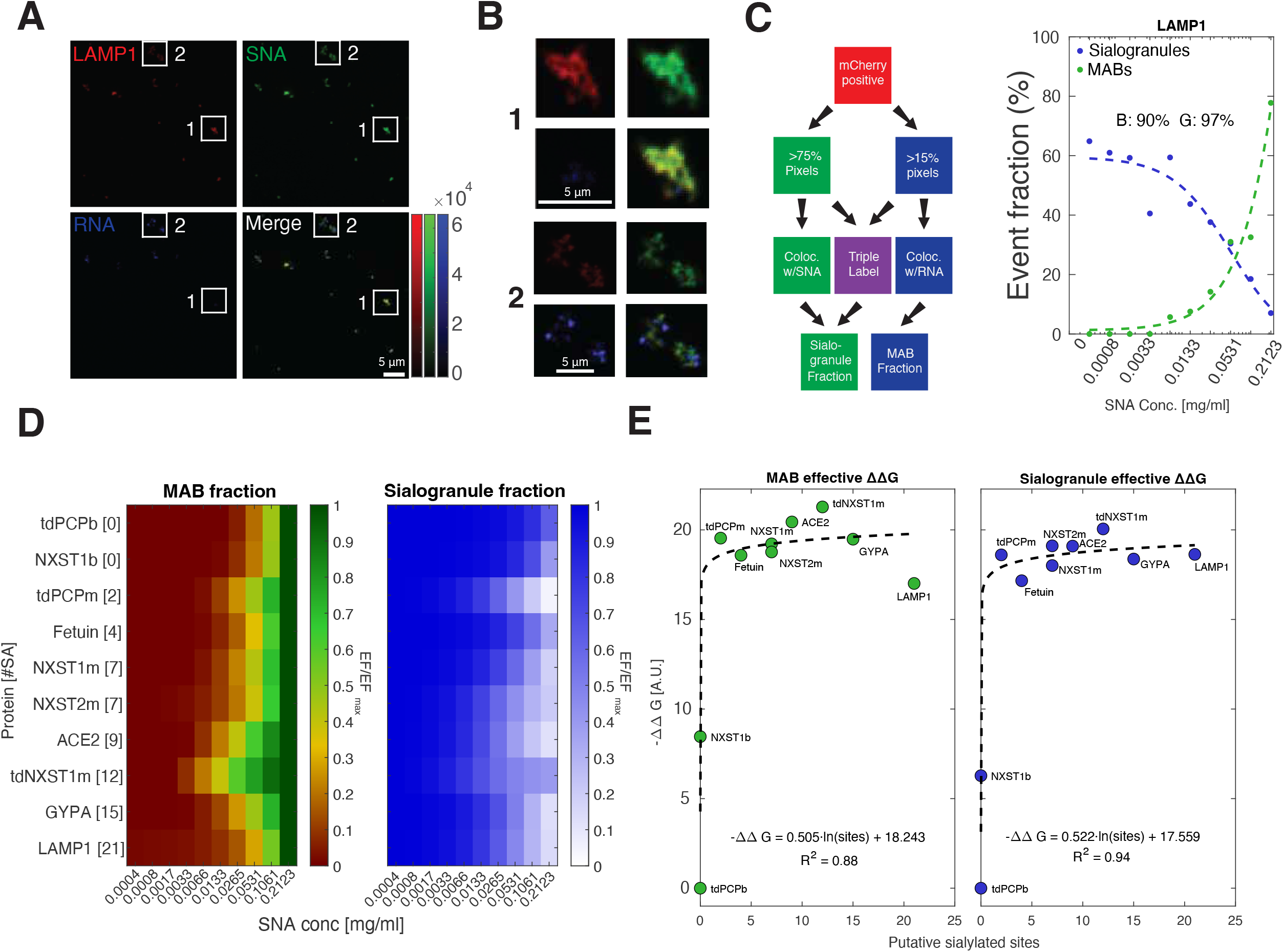
SNA-based MAB formation depends on the number of sialylated sites. **(A)** Typical field-of-view of SNA-LAMP1 MABs divided into individual channels. (Top-left) protein (mCherry). (Top-right) SNA (AF488). (Bottom-left) slncRNA (AF405). (Bottom-right) Merged panel with all three images overlayed. Scalebar: 5µm. **(B)** Close-up on the two features highlighted in panel A displayed via the same three separated channels and merged image. Scalebar: 5μm. **(C)** Left: diagram of event classification: a cluster of mCherry-positive events is colocalized with clusters of FITC- or AF405-positive pixels if the overlap is at least 75% and 15%, respectively. Triple labelled events are counted towards the SNA-binding bin. Right: LAMP1 dose-response curves (right) for sialogranule (blue) and MAB (green) event frequency produced based on the decision diagram. Percentages represent the goodness of fit to a Hill curve (B – blue, G – green). The dose-response curves for the other proteins is shown in Figure S3. **(D)** Dose responses (see Figure S3B for all proteins) displayed as a normalized heatmap for SNA binding (top) and RNA displacement (bottom). Each row was normalized by its maximal value. **(E)** The change in Gibbs’ free energy (ΔΔ*G*) computed based on the event frequency *K*_*d*_ (see methods) for both the granules and MABs is dependent on the number of putative sialylated sites. A logarithmic fit is demonstrated between the number of putative sialylated sites and either the accumulation MAB (left, R^2^=0.88) or the reduction of sialogranule events (right, R^2^=0.94).

The analysis shows (Figure 3C, dose-response, and Figure S3A) that for almost every putative sialylated protein, the fraction of MABs increases as a function of increasing SNA concentration, while simultaneously the frequency of sialogranules declines. The mean event frequency (EF) of each colocalization type were then fitted as increasing or decreasing Hill functions for either the MABs or sialogranules, respectively (see curves in Figure 3C and Figure S3B). To compare between the different candidates, each fitted event frequency was normalized by the maximal event frequency of the respective sialoprotein (Figure 3D). For MABs, the results show a dependency on the number of putative sialylated sites, where a significant FITC signal could be detected at lower SNA concentrations for a sufficient (n > 7) putative number of sialylated sites (Figure 3D, top). For sialogranule frequency, a gradual decrease in AF405 signal was observed as a function of increasing SNA concentration increases, as expected. Lastly, we extracted Hill constants (*K*_*d*_) from the fitted curves for each of the sialoprotein candidates, and normalized by the *K*_*d*_ for tdPCPb as reference, to estimate an effective avidity Gibbs free energy for SNA-granule biocondensate structure formation (*ΔΔG*, see Methods). To do so, we used the *K*_*d*_ extracted for both the increase in MABs, and the depletion sialogranules. For both cases, *ΔΔG* values exhibited a similar logarithmic dependency on the number of possible sialylated sites. This logarithmic behavior is consistent with predictions of a thermodynamic model (Figure 3D – fit, see Methods) that takes into account the sialic-acid valency of the sialoprotein. Thus, increasing the number of sialic acid modifications generates diminishing returns in terms of the gain in *ΔΔG* beyond a certain valency. Altogether, the SNA dose-response analysis provides further support that SNA-mediated phase transitions can differentiate between sialylated and unsialylated granules, and in addition also suggests that an estimate of the number of average sialylated sites on a target protein may be obtained from this avidity-based phase-separation assay.

### Masking or removing sialic acid greatly disrupts SNA binding and slncRNA displacement

To validate our phase-separation assay for the screening of highly sialylated granules, we wanted to test whether inhibiting SNA binding results in the restoration of granule events at the expanse of the frequency of MAB events. To do so, we opted to remove part or all of the N-linked glycans on LAMP1 or remove its sialic acids (Figure 4A). LAMP1 is known to carry both sialylated and unsialylated NLG patterns^37^. Endo H is a glycosidase that removes N-linked glycans that are not heavily sialylated^38,39^ (Figure 4A, black arrows), and treatment of Endo H on LAMP1 resulted in a polyacrylamide gel electrophoresis (PAGE) band shift of ∼20 kDa in LAMP1 bands (Figure 4B), similar to what is reported elsewhere^37^. This shift was evident for the heavily glycosylated main band (Figure 4B, black arrows) and for the secondary under-glycosylated isoform (Figure 4B, gray arrows). We also treated LAMP1 with a monomeric IFV’s neuraminidase (NA), a sialidase that cleaves sialic acid in any confirmation^40,41^ (including α2,6-SA) (Figure 4A, blue arrows). A PAGE assay revealed that NA-treated samples generate a broader smeared band as compared with the non-treated control (Figure 4C).

**Figure 4.**
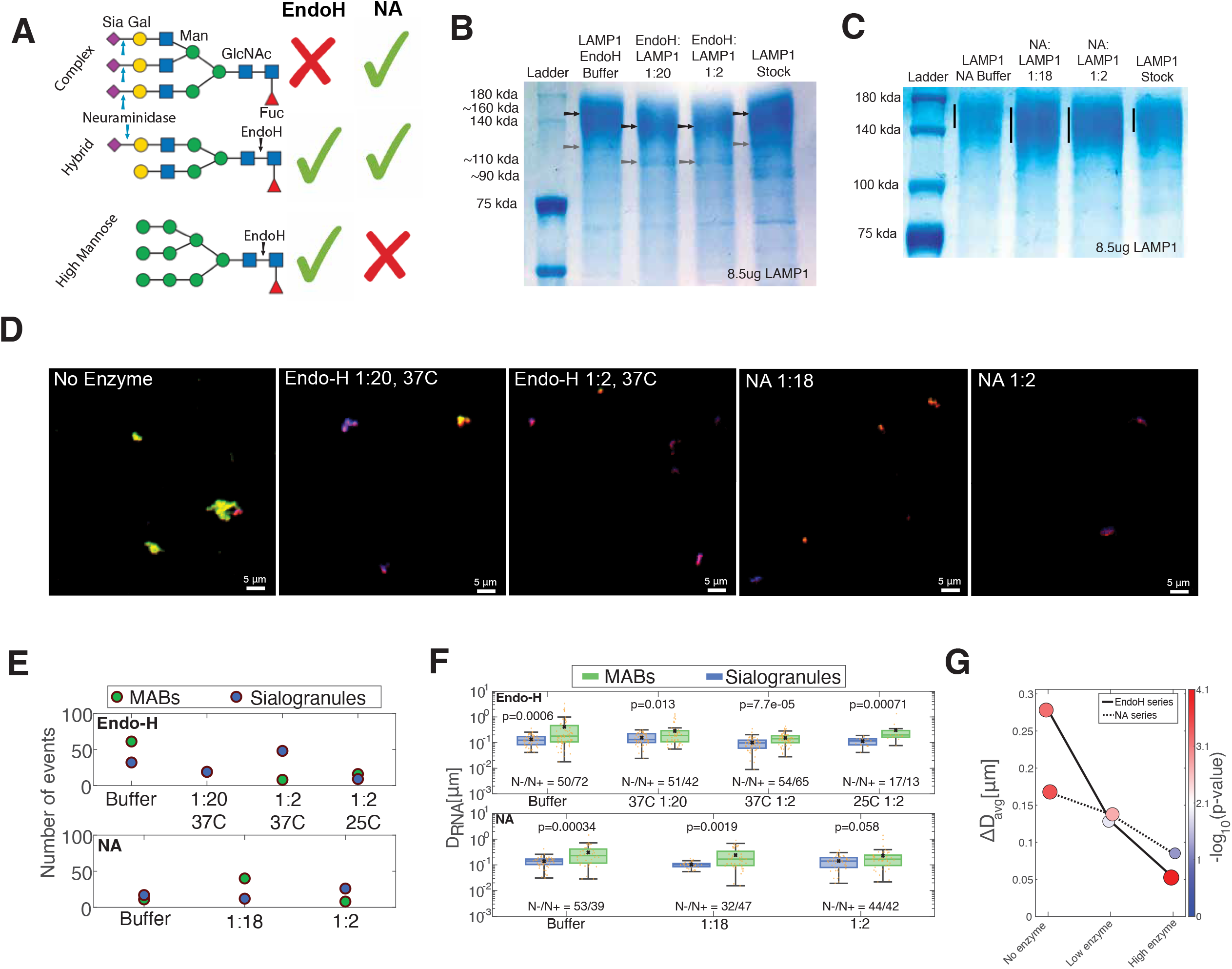
Disruption of SNA binding to LAMP1 by Endo H and neuraminidase treatments. **(A)** Schematic of the three main NLG patterns. Arrows mark where Endo H (black) and NA (blue) cleave. **(B+C)** PAGE results for Endo H **(B)** and NA **(C)**. Endo H treatment resulted in a band shift of ∼20kDa for both primary (black arrows) and secondary (gray arrows) of LAMP1, while NA treatment resulted in elongating (vertical arrows) and lowering the smeared main band. **(D)** Representative fields of view from different enzymatic treatments. Scale bar: 5 μm. **(E)** The number of events that are either MAB-like or granule-like according to the same parameters of Figure 3C. **(F)** *D*_*RNA*_ distributions for different conditions of Endo H (top) and NA (bottom) treatments. Blue and green boxes represent samples that were not incubated or incubated with SNA, respectively. A student’s t-test was performed for statistical analysis. **(G)** Δ*D*_*RNA*_ changes based on enzyme amounts for either Endo H- or NA-treated LAMP1 granules. Size and color represent the statistical significance reported in Figure 4F.

We then formed sRNP granules with Endo H- and NA-treated LAMP1 proteins and incubated them with SNA (Figure 4D). The results show that enzyme-treated LAMP1 formed sRNP granules that for the most part did not colocalize with SNA, while non-treated LAMP1 formed the usual MABs, as expected (Figure 4D – left). Moreover, treatment with increasing concentrations of Endo H (Figure 4D – second and third images, from left) or NA (Figure 4D – fourth and fifth images, from left) resulted in a recovery of granule event frequency in the presence of SNA (Figure 4E – blue circles) at the expense of the MABs that have become more sparse (Figure 4E – green circles). This contrasts with non-treated samples, where the number of MABs substantially exceeded the number of granules within the samples (Figure 4E – buffer). We also examined LAMP1 treated with Endo H at 25ºC, and our results show a more equal proportion of MABs and granules, consistent with the manufacturer’s reported catalytic efficiency of 40% for Endo H at that temperature (Figure 4E – top-right and S4).

Finally, to validate our analysis approach we measured for each condition the *D*_*RNA*_ values (as specified for Fig. 2) for all biomolecular condensation events that exhibited a distinct mCherry signature (Figure 4F) for the cases of no SNA and with SNA in the granule solution, respectively. Our results (Figure 4G) show, similarly to Fig. 2C, differences in *D*_*RNA*_ distributions that vary between weakly significant to non-significant between granules incubated without (blue) and with (green) SNA, respectively. This is consistent with our expectations as some of the granules even in the strongly treated samples (e.g. Endo H 1:2) still exhibit a small proportion of diffuse MABs. We then calculated Δ*D*_*RNA*_ for each condition by subtracting the mean *D*_*RNA*_ of the blue populations (no SNA) from the mean *D*_*RNA*_ of the green populations (with SNA). Our results show (Figure 4G) that Δ*D*_*RNA*_ decreased dramatically for Endo H-treated LAMP1 (∼0.05 μm). from non-treated LAMP1 (∼0.3 μm), amounting to a ∼81% reduction. When examining the correlation relationship between Δ*D*_*RNA*_ and the number of sialic acid modifications plotted for sialylated proteins in Fig. 2D, this result translates to a reduction to ∼3 available sialylation sites remaining on the Endo H-treated LAMP1. By contrast, for the NA-treated LAMP1 a more modest reduction in Δ*D*_*RNA*_ is obtained (∼49%), which translates to ∼6-9 available sites remaining on the NA-treated LAMP1. Taken collectively, these results confirm both an anti-glycan and more specifically anti-sialic acid activity by these enzymes, thus validating our sialogranules as sensors for sialic acid modifications.

### Decoy granules inhibit influenza infection using a pinball-like disrupted diffusion mechanism

As a final validation for our sialogranule screening pipeline, we proceeded to experimentally test LAMP1 candidate decoy granules for antiIFV functionality using an *ex vivo* IFV infection model^42^. Different amounts of the human IFV A/Puerto Rico/8/1934 strain (quantified by volume of viral stock mixed with the treatment and growth media) were used to infect human lung A549 cells in the presence of Endo H-desialylated LAMP1 granules, native LAMP1 candidate decoy sialogranules, or in the absence of any granules (Figure 5A, see Methods). 48h post-infection, cells were stained for viability and the presence of HA and measured using flow cytometry. Cells that were negative for viability dye (live cells) and positive for HA presence (infected) were used for the analysis (Figure S5A). The results show (Figure 5B) that the percentage of infected cells as a function of viral dose is shifted to the right (i.e for higher values) for the LAMP1 sialogranule treatment (green), as compared with the untreated (red) and Endo H-desialylated LAMP1 granule controls (blue). We then computed IC_50_ values for each treatment by fitting the virus dose response curve to a Hill function of order 1 (see Methods). The results show (Figure 5C) that the IC_50_ value obtained for the LAMP1 sialogranules increases by ∼2.25-fold, as compared with the IC_50_ values obtained for the untreated and the Endo H-treated desialylated LAMP1 granule treatment. This indicates that LAMP1 sialogranules necessitate a higher level of viral titer to yield a similar level of infected A549 cells, as the one observed for the controls.

**Figure 5.**
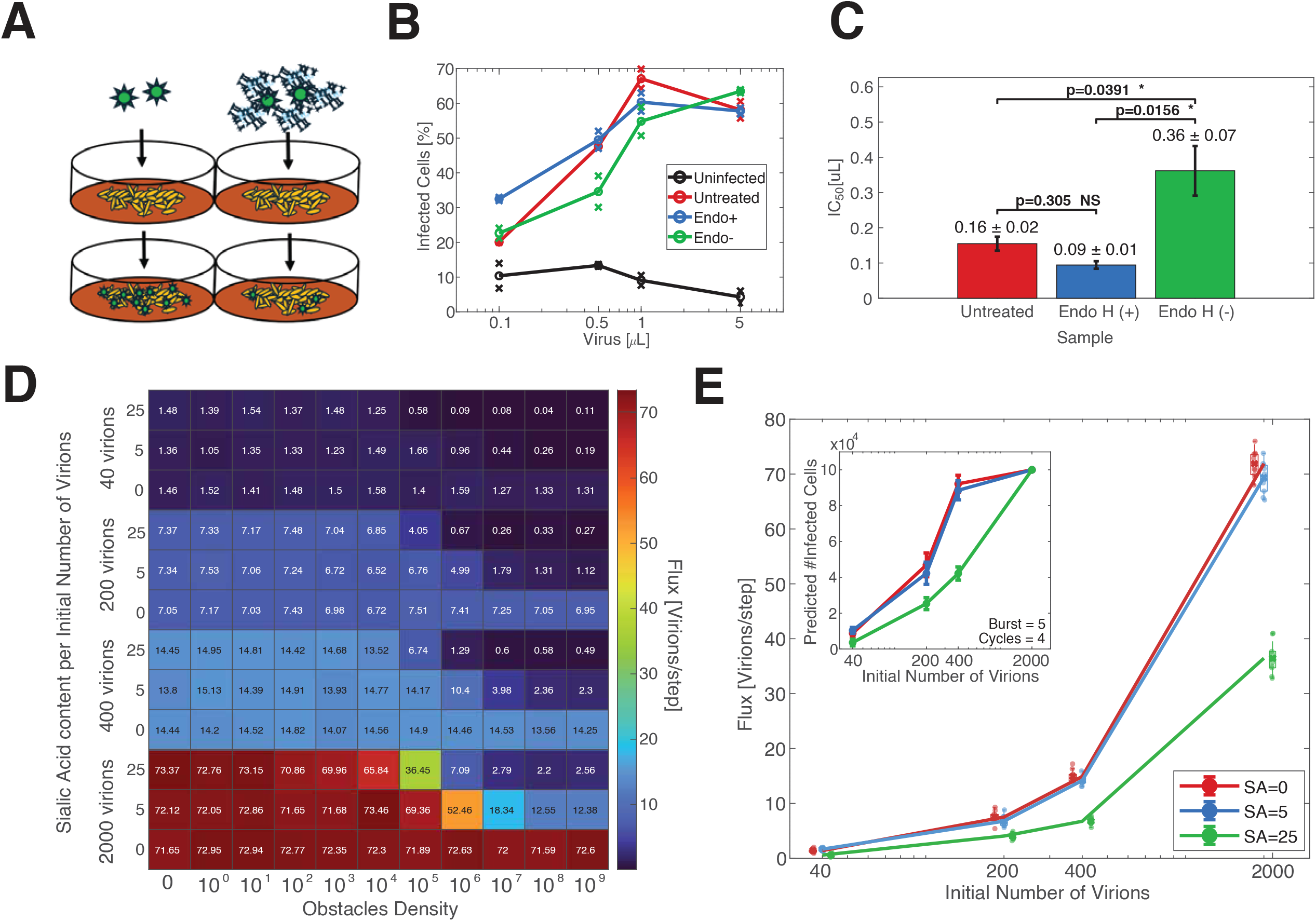
A disrupted diffusion model accounts for inhibition of IFV infection by LAMP1 decoy particles in A549 cells. **(A)** schematic of viral entry assay. Influenza virions (green objects) infected A549 cells (yellow objects) in the absence (left) or presence (right) of sialogranules. **(B)** Entry assay results for Endo H-untreated LAMP1 decoys (Endo-, green), Endo H-treated LAMP1 decoys (Endo+, blue), and no decoys (Untreated, red) for four increasing viral volumes. Uninfected cells (black) demonstrate false-positive binding of AF488-labelled anti-HA antibody (mean ∼8%). **(C)** Calculated mean IC_50_ values (half the volume needed for maximal level of infection). See Methods for calculation of error-bars and p-values. **(D)** Heatmap for the mean flux values of the simulated results in Figure S5B for different sialic acid content and number of virions. **(E)** Main: flux values and distributions from 100k obstacles density. Inset: trajectory of viral replication using the mean flux values for each simulated viral volume, assuming a burst size of 5 plaque-forming units (PFU) after 4 cycles of viral infection and replication. Error bars represent the standard deviation of propagated cells from the different runs of the simulation.

Finally, we wanted to estimate the extent of inhibition of IFV infection caused by the LAMP1 sialogranules given the observed shift in IC_50_. To do so, we invoked a simplified infection model (see Methods). In brief, our model assumes that a diffusion process^43^ controls the underlying dynamics of virions as they seek a cell to infect. Our decoy particles are ∼10-fold larger than typical ∼50-100nm diameter virions, and serve as partly static obstacles positioned at random locations in space, similar to how mucin proteins occupy the mucosal layer of the lung^44–46^ and slows IFV motility in such a medium^45–48^. The IFV HA affinity to α2,6 sialic acid is generally low^26^, and therefore receptor strength is represented by the extent of a sialoprotein candidate’s sialylation. In this simplified scheme, increasing the number of decoys and the extent of their sialylation leads to a disrupted diffusion process by extending the time a virion remains bound to a decoy granule. This, in turn, leads to a reduction in virion flux on the cell layer (Figure S5B), resulting in reduced infection. The model treats the decoy particles as obstacles in the Brownian motion of IFV virions, which are able to attach (using the HA protein) and detach catalytically (using the NA protein) ^44–46^.

We simulated the diffusion of various number of viral particles towards a cell layer through a dense field of obstacles with varying degrees of sialylation (0, 5, and 25, imitating non-sialylated granules, Endo H-desialylated LAMP1 granules, and native LAMP1 sialogranules, respectively). At the end of each run, the mean flux of virions per time step was plotted as function of obstacles density (Figures 5D and S5B). Increasing the decoy density resulted in a sharp drop of viral flux, with 10^5^ being the obstacle density for highly sialylated obstacles where the mean flux of virions on the cell layer is cut by half as compared with that of 0 sialic acids (Figure 5D and S5B, right), resembling the experimental results. We continued to plot the flux results of 10^5^ obstacles for each of the different virus amounts (Figure 5E). These values were used as initial values for a propagation simulation for each SA curve, reaching a limit of ten thousand cells (seeding density in the experimental system). We assumed a burst size of 5 plaque-forming-units (PFU), for each of the three sialic acid contents (Figure S5C). Low or null sialic acid curves reached the threshold rapidly (just after flux values of 400 virions), while the curve for highly sialylated obstacles only reached cell saturation for the flux value of the highest virion count (Figure 5E, inset), resembling the trends obtained in the experiment. This then allows us to estimate that the LAMP1 sialogranules’ reduced IFV infection by ∼50% (e.g. compare green value to blue and red for N=2000 and 400 virions in Figure 5E), consistent with the IC_50_ results.

## Discussion

In this work, we demonstrated that sialogranules composed of RNA and sialylated proteins can function as high-avidity Velcro-like glycoNPs. Our sialogranules bind lectin targets with high avidity leading to a conformational change of the granular phase into a multiphasic agglutinated biocondensate (MAB). Detailed analysis of the MAB revealed that morphological features of the biocondensate correlate with the putative number of sialic acid modifications on the sialoproteins. As a result, we were able to empirically rank our sialogranules based on their sialic-acid content leading to the identification *in vitro* of the LAMP1 sialogranule as a functional candidate IFV decoy via entry experiments in human A549 cells.

A deeper examination of our results allows us to draw wider conclusions. The transition into the MAB state and its logarithmic dependence on the number or density of sialic acid modifications on the sialogranule (Figures 2D, 3E, and S3) is consistent with a simple avidity-based kinetic binding model (see Methods) that provides a biophysical mechanism for the regulation of biomolecular condensates by ligands. As a result, this regulatory effect should be generalizable to other post-translational modifications, and the structural melting of granules can form the basis of a new type of “avidity-binding” assay. Such an assay can be employed with relative ease and could serve as an alternative to existing nanoparticle-based, ELISA-based, or mass spectrometry-based diagnostic approaches. In particular, our ability to differentiate between two genetically identical protein products based solely on the presence of post-translational sialylations due to the cell-type in which they were expressed (Figure 1D) provides a simple proof-of-concept as to the validity and utility of this assay.

Furthermore, as part of the validation process, we found that the glycosidases Endo H and NA exhibited apparent catalytic activity via gel-shift assays and our sialogranule screen. While these results for NA were expected, the apparent activity of Endo H was a surprise, since there have not been other reported instances alluding to an Endo H activity on sialylated glycoproteins. Given the additional validation for Endo H activity that we observed via the viral entry assay, we postulate that either Endo H has substantial sialidase activity contrary to previous reports^38,39^, or it has a binding affinity to sialylated glycans and is able to mask them from SNA and HA interaction but not necessarily remove them. Either way, EndoH generated a substantial reduction in sialogranule avidity to lectins inhibiting both the transition to the SNA-based MAB state and the ability of the granules to prevent IFV infection. Consequently, our sialogranules constitute a new class of diagnostic glycoNPs, which can open new avenues of research into characterizing the amount of post-translational modifications on a given protein, and the various catalytic effects and/or binding capacity of glycosidases.

Finally, the viral entry assay results for IFV imply that the proposed platform may be used as a basis for high-avidity glycan-lectin therapeutics. The proposed Velcro-like therapeutic will comprise of a slncRNA cassette and potentially any number of slncRNA-binding soluble receptors or glycanated antiviral peptides. By incorporating different RNA binding protein moieties and their respective binding sites, this platform can, in theory, be programmed to inhibit infection of single or different combinations of virus species, address bacterial pathogenicity, and possibly aid in cancer therapy. Consequently, our platform provides a pathway for the development of new therapeutic strategies that take advantage of other low-affinity ligand-target interactions.

### Concluding Remarks

Synthetic sialogranules represent a programmable class of genetically encodable glyco-nanoparticles that integrate multivalent glycan sensing with functional antiviral activity. By converting weak lectin-glycan interactions into an avidity-driven phase transition, we establish a quantitative, morphology-based detection platform in which multivalent ligand binding remodels granule architecture. Importantly, the granule platform self-assembles from a mix of synthetic long non-coding RNA scaffold and glycosylated proteins and is therefore capable of incorporating unlimited glycan complexities dictated by mammalian biosynthesis. As a result, glycogranule-based assays might offer a generalizable strategy for measuring post-translational modifications and molecular interactions, without relying on surface immobilization or conventional nanoparticle chemistry. From a therapeutic perspective, the LAMP1 sialogranule antiIFV activity and the ACE2-decoy granule antiSARS2 activity indicate that the granule platform can be a modular antiviral platform with a potential for broad-spectrum efficacy. Consequently, this combination of simplicity and biochemical richness opens a path toward customizable, high-avidity glycan nanoparticles for a wide variety of diagnostic and therapeutic applications.

### Outstanding Questions

- **How generalizable is granule-based detection?** Can avidity-based phase-transition signatures reliably distinguish other post-translational modifications?
- **Can phase-separation-based nanoparticles be used to detect and quantify other weak specific molecular interactions?** Beyond sialylation and ACE2-RBD binding, can this platform serve as a universal biophysical assay for weak ligand-receptor interactions and enzyme-substrate kinetics?
- **Can multi-receptor granules achieve true broad-spectrum activity?** Is it feasible to engineer a single granule displaying multiple soluble receptors (e.g., ACE2, LAMP1, heparan sulfate-binding motifs) to inhibit unrelated virus families simultaneously?
- **Can a glycangranule or lectingranule platform be expanded to treat other maladies?** Can a granule platform be used in cancer therapy and to inhibit pathogenic bacteria?

### Technology Readiness

The synthetic RNA-protein (sRNP) granule platform is currently positioned at early Technology Readiness Level (TRL) 4, reflecting validation of the core technology in a controlled laboratory environment with functional performance in a biologically relevant human cell system. In this study, we demonstrate mechanistic and quantitative validation of avidity-driven phase transitions, showing that SNA-induced multiphasic agglutinated biocondensate (MAB) formation scales with predicted sialylation valency and permits extraction of effective dissociation constants and ΔΔG values. Enzymatic perturbation using Endo H and neuraminidase confirms glycan-dependent specificity and reversibility of the phase-transition readout, while dose-response analysis establishes reproducible avidity-dependent behavior across multiple constructs. Most importantly, LAMP1-based sialogranules produce a ∼2.25-fold shift in IC50 and ∼50% reduction in influenza A viral entry in A549 human lung epithelial cells, supported by a predictive diffusion–attachment model that links sialylation density to antiviral efficacy.

Although these results demonstrate modularity, quantitative mechanistic control, and functional antiviral activity in a relevant epithelial infection assay, the platform has not yet been evaluated in complex physiological environments or in vivo systems. Advancement toward higher TRLs will require validation in airway-mimicking mucus or serum, testing in organoid or small-animal infection models, and systematic assessment of pharmacokinetics, stability, biodegradation, and immunogenicity. Establishing scalable and reproducible production workflows for glycosylated decoy proteins will further determine translational feasibility. Collectively, the current data support classification as early TRL 4, with a clear pathway toward preclinical development.

## STAR⍰Methods

### Key resource table

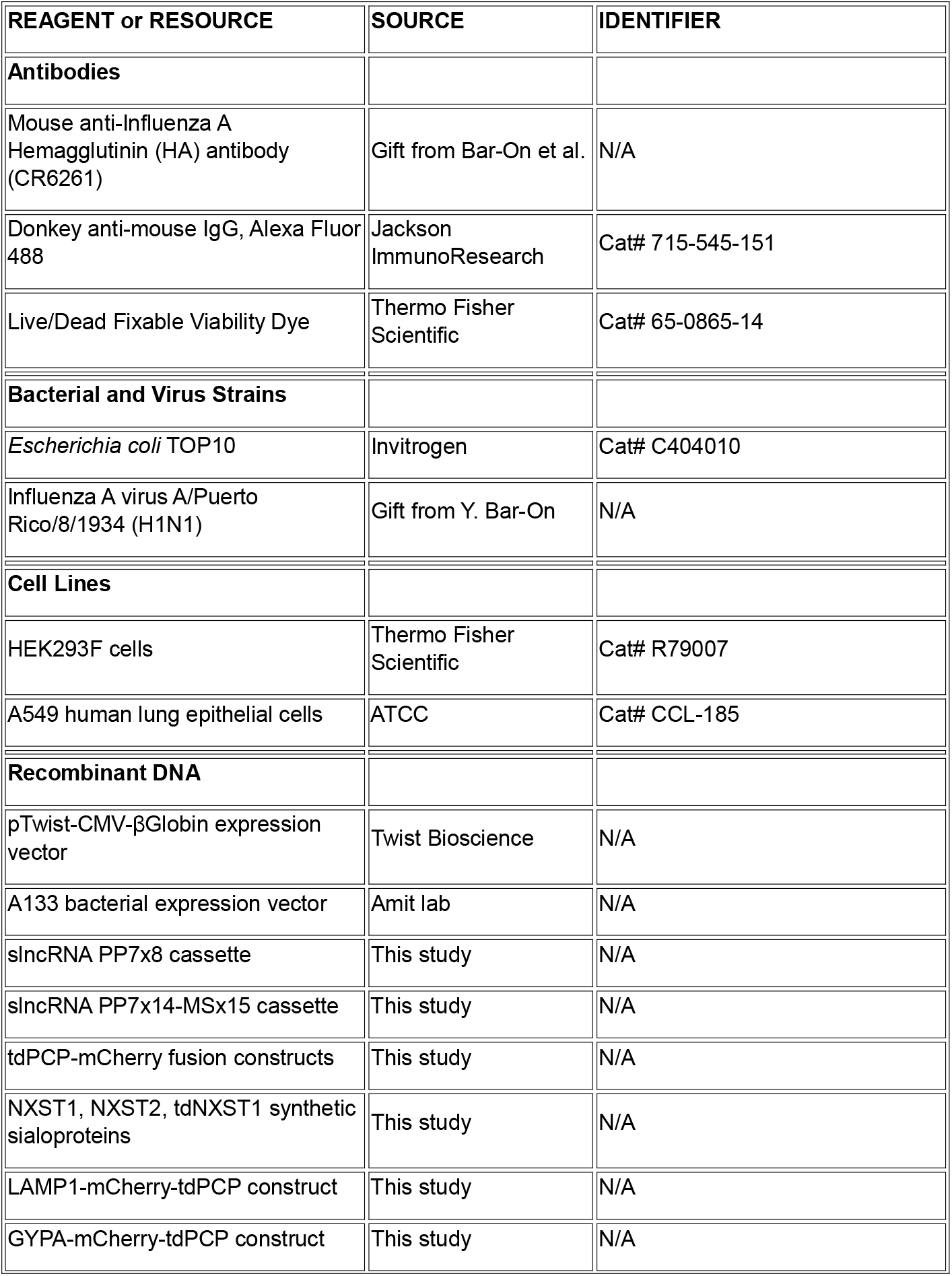

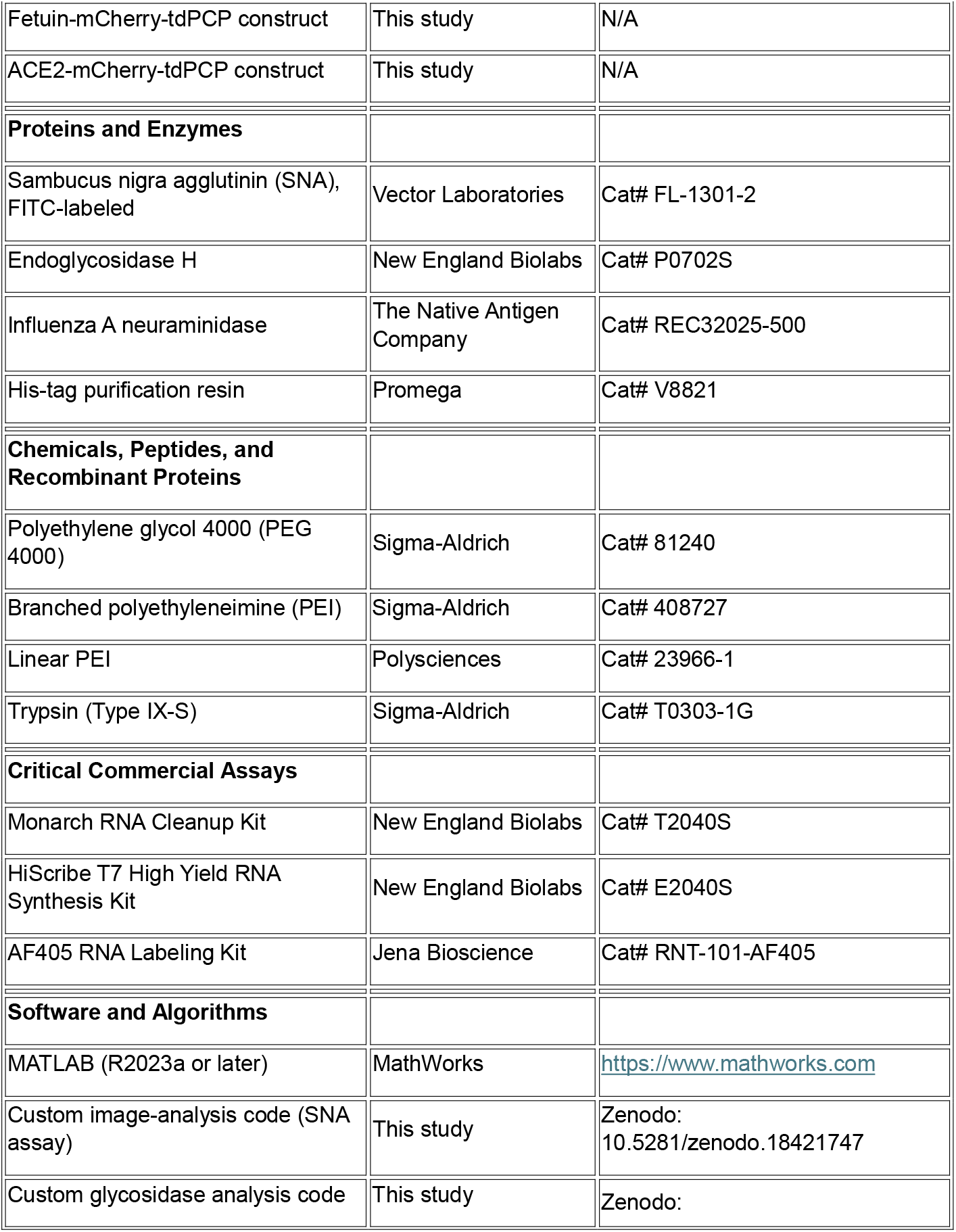

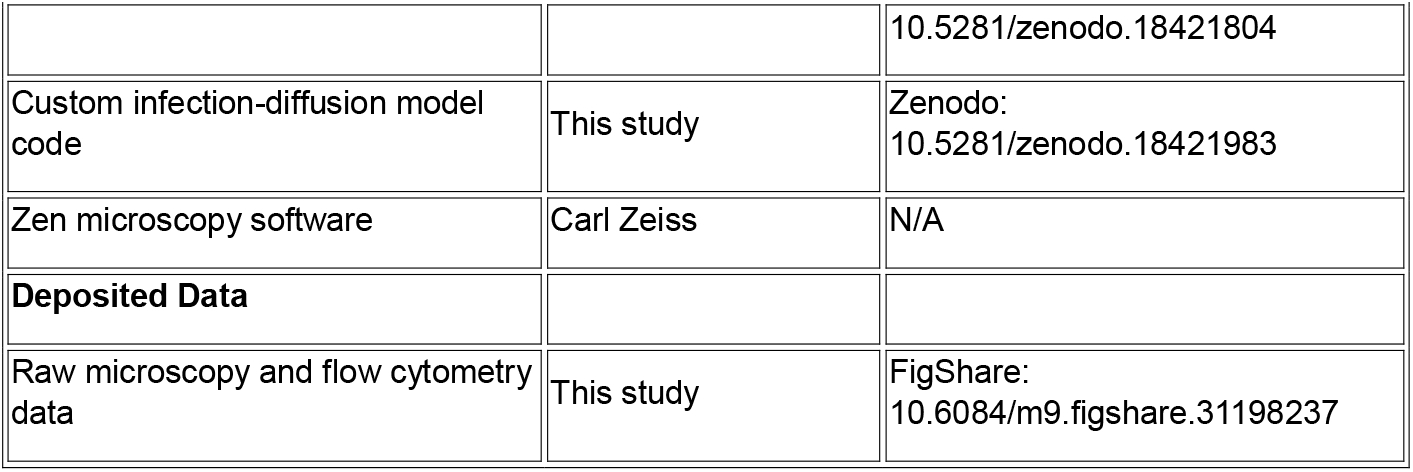

### Plasmid construction

Protein genes were ordered as either Gene Fragments (Twist Bioscience) or gBlocks (Integrated DNA Technologies, IDT). For the natural proteins LAMP1, GYPA, Fetuin/AHSG and ACE2, the sequence that was cloned was extracted from UniProt and contained the protein’s native signal peptide and the extracellular domain, namely amino acids 1-382, 1-91, 1-367, and 1-740, respectively. For the NXST peptide variations, the signal peptide of ACE2 was used. The NXST peptide is comprised of five alternating repeats of the known sequon for N-linked glycosylation N-X- (S/T) with six amino acids separating between each sequon. For the mammalian tdPCP protein, the signal peptide of ACE2 was used directly upstream to the protein sequence.

All but two proteins were expressed in HEK293F cells. All of the proteins produced in mammalian cells were made from genes cloned into a pTwist-CMV-BetaGlobin vector downstream of the β-globin intron. An affinity tag comprised of six histidines (his-tag) for purification purposes was positioned immediately upstream to the stop codon of all coding sequences. Bacterial tdPCP (tdPCPb) was cloned into A133 plasmid under N-butyryl-L-Homoserine lactone (C4HSL) induction and expressed in TOP10 *E. coli* cells. NXST1b was cloned into A133 plasmid using the coding region from the pTwist-CMV-BetaGlobin vector and expressed in the same manner as tdPCPb. Plasmids containing the synthetic long non-coding RNA (slncRNA) cassettes were created as previously described^20^.

### In vitro transcription and purification of RNA cassettes

RNA transcription of synthetic RNA cassettes was done as previously described^20^. Briefly, a vector containing the slncRNA DNA sequence downstream of a T7 promoter, flanked by two EcoRI restriction sites, was digested with EcoRI-HF (NEB, cat. R3101L) per the manufacturer’s instructions to form a linear fragment. The restriction enzyme was then heat-inactivated by incubating the restriction reaction at 65□°C for 20□min. For fluorescently labeled RNA, 1µg of the restriction product was used as template for in vitro transcription using HighYield T7 Alexafluor-405 (AF405) RNA labeling kit (Jena Bioscience, cat. RNT-101-AF405), according to the manufacturer’s instructions. Non-fluorescent RNA was transcribed using the HiScribe™ T7 High Yield RNA Synthesis Kit (NEB, cat. E2040S). Following in vitro transcription by either kit, the reaction was diluted to 90□µl and was supplemented with 10□µl DNAse I buffer and 2µl DNase I enzyme (NEB, cat. M0303S) and incubated for 15□min at 37□°C to degrade the DNA template. RNA products were purified using Monarch RNA Cleanup Kit (NEB, cat T2040S) and stored in −80°C.

### Cell maintenance

A cryotube with 1ml 10%v/v DMSO (Sigma-Aldrich, cat. D2650) of 10^7^ HEK293F cells/ml (Thermo Fisher, cat. R79007) was thawed as follows: the cryotube’s content was transferred into 4ml fresh Freestyle media (Thermo Fisher, cat. 12338018), prewarmed to 37°C, and then immediately centrifuged at 500xg for 5min. Supernatant was removed, and cell pellet was resuspended in 7ml of fresh Freestyle media and transferred to 23ml of prewarmed Freestyle media in a 125ml flat-bottom flask (TriForest, cat. TF FPC0125S). Cells were then moved to an incubator and grown under 37°C, 8% CO_2_ and humid conditions (a in-hood basin with 500ml autoclaved DI water, supplemented with 1%v/v Aquaguard-1 solution (Sartorius, cat. 01-867-1B) before use). The cells were placed on an in-incubator orbital shaker which rotated at 135rpm. These conditions were in accordance with the cells’ manual.

The cell concentration exceeded 1.2×10^6^ cells/ml 3-4 days after thawing, at which point cells were passed by diluting them 1:5 in a final volume of 30ml fresh growth media. Cells were routinely passed every 3-4 days, as cell concentration was sufficient. Cells were checked for mycoplasma in accordance with manufacturer protocol (Vazyme, cat. D101-02).

### Cell transfection

24h before transfection, HEK239F cells were seeded at 0.6-0.7×10^6^ cells/ml in 30ml fresh Freestyle media and allowed to grow overnight. On the day of transfection, cells were diluted to 1×10^6^ cells/ml. Per flask containing 30ml of culture, up to 40µg of plasmid DNA was diluted in OptiMEM buffer (Thermo Fisher, cat. 31985070) to a final volume of 600µl. 120µl of 0.5mg/ml branched PEI solution (Sigma-Aldrich, cat. 408727) or 60µl of 1mg/ml linear PEI (Polysciences, cat. 23966-1) was diluted in OptiMEM buffer to a final volume of 600µl. The PEI-OptiMEM solution was added to the DNA-OptiMEM solution (final mixed volume of 1.2ml) and incubated at room temperature for 15min. Each flask was then transfected with the appropriate plasmid within 5 minutes. Transfected cells were maintained for 5-7 days, during which time the growth media changed color to red due to the mCherry label in the recombinant proteins.

### Protein extraction from HEK293F cells

1ml of nickel (Ni)-coated beads (Promega, cat. V8821) were transferred into 50ml conical tubes and were allowed to settle. 400µl of supernatant was removed and beads were washed in 2.5ml of equilibration buffer (50mM NaH_2_PO_4_, 300mM NaCl, 5mM imidazole, pH 8). After the beads resettled, 2ml of the supernatant was removed. Transfected cells were centrifuged 5-7 days post transfection at 5000rpm for 20min at 4°C. Up to 40 ml supernatant was transferred to a 50ml conical tube with settled beads while cell pellet was discarded. Bead-protein mix was incubated at room temperature for 1h in an end-over shaker. After incubation, the solution was transferred to a gravity column (Bio-rad, cat. 731150) and ∼100µl of the flowthrough was collected. Next, beads were washed three times using 5ml of wash solution (50mM NaH_2_PO_4_, 300mM NaCl, 20mM imidazole, pH 8), and ∼100µl was collected from the first wash. Finally, beads were resuspended inside the column in 2ml elution buffer (50mM NaH_2_PO_4_, 300mM NaCl, 500mM imidazole, pH 8) and left to incubate for 15min, after which the entire elution volume was collected in 2-3 fractions.

### Protein extraction from TOP10 *E. coli* cells

TOP10 *E. coli* cells (Invitrogen, cat. C404010) were grown in 5ml Luria broth (LB), supplemented with 100µg/ml ampicillin (Sigma-Aldrich, cat. A9518), at 37°C and 250rpm overnight. The following day, the starter content was transferred to 500ml TB (24g yeast extract, 20g tryptone, 4ml glycerol, in 900ml DI water, autoclaved, then added 100ml of 10x phosphate buffer to final concentrations of 17mM KH_2_PO_4_ and 72mM K_2_HPO_4_), supplemented with 100µg/ml ampicillin and 100µM C4HSL (Cayman Chemical, cat. 10007898), and grown overnight at 37°C and 250rpm. The next day, cells were allowed to grow until reaching an optical density of >0.45, before being centrifuged at 6000rpm for 10min. Supernatant was removed, and cell pellet was resuspended in 30ml of resuspension buffer (50mM Tris base pH=8, 100mM NaCl, 0.02% sodium azide, in 1L DI water, titrated to pH=7), before being passed four times through an EmulsiFlex-C3 homogenizer (Avestin Inc.). Finally, cellular debris was centrifuged at 13000rpm for 30min at 4°C and supernatant was transferred into a new 50ml conical tube. Purification of proteins was executed as described for proteins produced in HEK-293F cells, after adding up to 40ml clarified cell lysate to settled Ni-coated beads.

### Verification of extracted proteins using SDS-PAGE

15µl of each fraction (flowthrough, wash, elution) per protein was taken to a sodium dodecyl sulfate polyacrylamide gel electrophoresis (denaturing SDS-PAGE), using a 10% acrylamide concentration in the resolving gel. 0.1mg/ml of purified BSA (NEB, cat. B9000) was used as positive control. Samples were boiled at 95°C for 10min before loading onto the gel. Gel running conditions were 150V for 70min. Afterwards, the gel was stained using Instant Coomassie Blue (Abcam, cat. Ab119211) for 15min.

### Buffer exchange of extracted proteins

The high imidazole content from the elution buffer was diluted ∼1:10^4^ by using an Amicon filtration tube (Millipore, cat. UFC200234) according to the manufacturer’s protocol. In brief, PBS×10 was added to the eluted protein samples as 1/9 volume equivalent its post-extraction volume. Then, PBSx1 was added to dilute the imidazole concentration to 100mM, the maximal concentration recommended by the manufacturer’s protocol. The diluted protein solution was loaded onto the Amicon tube in multiple loading steps and centrifuged for 15min at 5000rpm several times, until reaching 400-500µl retentate volume. In between each centrifuge, protein precipitate was resuspended by carefully pipetting inside the column. Once arriving at the appropriate volume range, 500µl of PBSx1 was added to the protein solution inside the column, diluting the imidazole ∼2-fold. This process was repeated eleven times to lower the imidazole concentration to approximately 50µM.

### Formation of RNA-protein granules

For microscopy assays, 250fmol slncRNA PP7×8 was mixed with any of the proteins at a molar ratio of 100:1 protein-to-RNA molecules. For agglutination assays and viral entry assays, proteins and slncRNA were incubated in 7µl of granule solution (750mM NaCl, 1mM MgCl_2_, 10% PEG 4000) and completed with water to a final volume of 21µl per reaction. The mixture was incubated at room temperature for 1h before being taken for imaging. For viral entry assays, the same protocol was maintained with a protein:slncRNA ratio was 10:1 (102.6pmol slncRNA), and the slncRNA was PP7×14-MSx15 cassette.

### Confocal microscopy imaging

For all imaging, we used an LSM-700 (Carl Zeiss) confocal microscope, equipped with a BIG (GaAsP detector) unit, using a Plan-Apochromat 63x/1.40 oil objective. Images were captured using Zen imaging software (version 8.0, 2012). A pinhole of 1.5µm was set for each laser. Emission windows were 578-800, 300-583, and 300-483 for the 555nm (mCherry), 488nm (FITC), and 405nm (AF405) channels, respectively, for all assays. For each assay we used standard microscopy slides with 1.5H coverslips, and each sample was imaged using 5µl.

### Confocal agglutination assay of RNA-protein granules with SNA lectin

2mg/ml stock concentration of FITC-labelled SNA Lectin (Vector Labs, cat. FL-1301-2) was serially diluted 2-fold nine times in biological water. 1.9µl of the appropriate SNA lectin dilution was added to 16µl of granules and incubated for 10min at room temperature. Samples were then imaged. Proteins were imaged with or without RNA. Laser intensities were 2% for all dyes. Gain values were 650, 550, 750 for mCherry (555nm), FITC (488nm), and AF405 (405nm), respectively. To optimize time, an averaging value of 4 and a depth of 16bit were set. Repeats were taken on separate days, each time acquiring between three to seven fields of view per sample. All fields of view were analyzed using MATLAB code written for the analysis (see code availability statement).

### Morphological analysis of agglutination assay

mCherry-positive events were defined as a cluster of 8 or more pixels that have an mCherry channel intensity value higher than a selected per-protein threshold (see code). Sialogranules-only samples (i.e. without SNA) had a generally lower mCherry-threshold than SNA-containing samples, as SNA agglutination concentrated the mCherry signal. FITC- and AF405-positive pixels were ones with an intensity value greater than global selected thresholds for each channel, irrespective of the bounded sialoprotein.

The center of gravity (CoG) of clusters of mCherry-labelled events was calculated, and the CoG of AF405-labelled pixels that occupied the same space as mCherry-positive events was similarly calculated. Then, the distance *D*_*RNA*_ between AF405 CoGs and mCherry CoGs for the mCherry-labelled cluster of pixels was calculated. The calculated distances in the three lowest and three highest SNA concentrations were pooled (Δ*D*_*RNA*_), and the means distances distribution were analyzed for statistical significance using student t-test.

### SNA lectin dose-response analysis

The thresholds used for this analysis were the same as the ones used for the confocal agglutination assay analysis. mCherry-positive events were assigned to one of four colocalization groups: (1) colocalization of mCherry, FITC and AF405 signals, (2) colocalization of mCherry and AF405 but not FITC, (3) colocalization of mCherry and FITC but not AF405, and (4) no colocalization. Assignment into the four groups was done by calculating how many of the mCherry-positive pixels were co-labelled with either FITC- or AF405-positive clusters, both, or neither. For FITC colocalization, 75% of an event’s pixels had to be FITC-positive. For AF405-positive events, mCherry-labelled events had to share at least 15% with AF405-positive events, as RNA was clustered in the periphery of mCherry-positive events as more SNA was added. The number of events in each of the four bins was divided by the total number of events per lectin concentration per protein, to yield the mean event frequency (EF) per bin per SNA concentration for each sialoprotein. Finally, triple-labelled events were counted towards the SNA-bound events, as SNA-sialoprotein colocalization was the predominant interaction for these events.

The error of the mean probability was calculated according to multinomial distribution approximation as follows (equation #1):

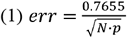

Where *err* is the error, *N* is the total number of observed events, and is the calculated probability per bin per lectin concentration per sialoprotein.

Analysis of the yielded dose response curve allowed for the extraction of dissociation constant (*K*_*d*_) values for either SNA binding or RNA exclusion. We used Matlab fit function for the following models (equations #2 and #3):

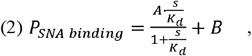

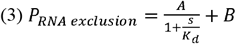

Where *A* is the amplitude of response between [baseline value, maximal value], *B* is the intercept for the dependent variable P, *K*_*d*_ is the dissociation constant, is the independent variable (SNA concentration), and *P* is the probability. SNA concentration was used as mg/ml, and therefore the *K*_*d*_ units are the same.

For the fit guess, *K*_*d*_ was set as the median probability per protein series.

Finally, the change in Gibbs free energy was calculated as follows (equation #4):

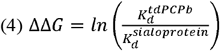

Where 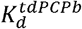 is the dissociation constant of the tdPCP-b protein produced in *E. coli*, which is not sialylated, 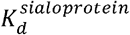 is the dissociation constant of a sialoprotein, is the natural logarithm, and ΔΔG is the change in Gibbs free energy.

### Numerical model for IFV virion flux measurement

We developed a numerical simulation to study a disrupted diffusion process of particles in a 3D bounded space with obstacles, incorporating particle attachment, hit counts, and particle generation mechanisms. The primary goal was to analyze the flux of particles hitting a cell layer, i.e., “the measurement wall”, arbitrarily designated as the x = 0 plane.

Model assumptions include (1) 3D bounded space with limits at [*L*_*x*_, *L*_*y*_, *L*_*z*_]; (2) particles undergo Brownian motion, characterized by a diffusion coefficient *D*; (3) virion flux is measured on the plane of *x* = 0; (4) the space contains spherical obstacles which simulate a phase-separated granules containing decoy receptors, treated statistically as a uniform spatial density *ρ* (see below); (5) upon hitting an obstacle, a virion becomes immobilized for a specified number of time steps *t*_*attach*_ before being released in a random direction, simulating interactions between IFV’s HA protein and a sialoreceptor; and (6) upon release, virions resume standard Brownian motion with no memory of previous direction, simulating the function of IFV’s neuraminidase (NA) protein.

The overall number of virion particles remained the same, and whenever a virion reached x = 0, a new virion was generated randomly in the 3D space. At the start of the simulation, *N* virion particles and *K* obstacles are initialized at random positions withing the bounded space. At each simulation step, particle positions are updated based on a diffusion process (equation #5):

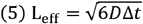

Where *D* represents the diffusion coefficient and Δ*t* is a time step.

Particles are reflected back into the bounded space if they hit a boundary, as follows for each coordinate *u* ∈ {*x,y,z*} with bounds [0,*L*_*u*_](equations #6-7):

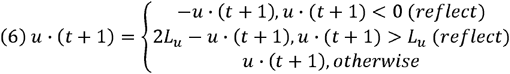

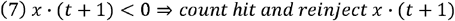

Viral diffusion takes place for all non-attached virions as a Brownian displacement:Δ_*x*_,Δ_*y*_,Δ_*z*_ ∼ *N*(0,2*D* · *dt*). Then the model computes the probability that a virion encounters an obstacle during a diffusion step through an obstacle density field (equations #8-10):

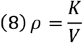

Where *ρ* is obstacle density, *K* is the number of obstacles, and is the 3D volume.

The number of collisions per unit of time is given as (equation #9):

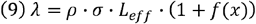

Where *λ* is the rate of collisions corresponding to the mean “on”-rate (i.e the rate by which the virions bind the obstacles), *σ* is the geometric cross-section of an obstacle, *L*_*eff*_ is the effective diffusion step’s length, given by equation #1, and *f(x)* is some function of the number of sialylated sites in a single protein within an obstacle. To a first approximation, it is customary to assume that *f(x)* is second order contribution and can thus be neglected. Importantly, to account for the heterogeneity in obstacles size (representing the varying size of our decoys, reported previously^22^ and also observed in this work), each obstacle was assigned an effective geometric cross-section *σ*, drawn uniformly from a range of 30 to 100 units at each update interval (every accumulated 5% of the simulation steps, representing content and size changes, reported previously^20^).

The probability of at least one encounter (*P*_*hit*_)is given as a Poisson distribution (equation #7):

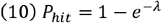

A numerical encounter corresponds to a binding event. This, in turn, yields a penalty of time steps corresponding to the mean “off”-rate, where the virion remains stationary (i.e. bound to the obstacle) for a preset subset of time-steps. Subsequently, diffusion continues in a randomized direction. In this simulation, the penalty or “off” rate is calculated thusly (equation #11):

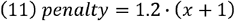

Where *x* is the number of sialylated sites in a single protein within an obstacle, and 1.2 is some constant. This constant is not derived, nor does it represent a physical quantity. Loosely, it can be thought of as a the off-rate for a non-specific collision process. The dependence on x is based on the following assumptions. Each sialogranule has a particular density of sialic acid modifications that are displayed for interaction. That density depends on the # of putative sialylation sites for each sialoprotein. Since the virion likely interacts with multiple sialylated sites, the “off”-rate must therefore depend on the sialylation density. Therefore, to a first approximation, we assume a linear dependence of the “off”-rate or penalty on the number of putative sialylation sites. Finally, the number of particles hitting the measurement wall is recorded, and the flux rate is calculated at the end of the simulation as (equation #13), where *S* is the total number of simulation steps (1000):

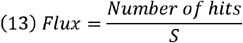

Note, that given the definitions of the mean “on”-rate (eqn.9) and “off”-rates (eqn. 11), it is possible to derive an expression for the effective binding affinity for the sialogranule interaction with a ligand of any sort as follows:

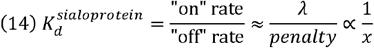

By plugging this term into equation #4, which provides a relationship between the change in Gibbs free energy for the SNA-sialogranule interaction as a function of a fitted effective binding affinity, we get:

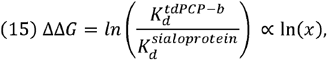

which is indeed supported by the data (see fits in Fig. S3B), providing further validity to our modelling approach.

### Endo H treatment of LAMP1

20µg of LAMP1 was incubated with 2µl of Endo H (NEB, Cat. P0702S) or a 10-fold dilution, corresponding to 1:2 and 1:20 enzyme:LAMP1 molar ratios, according to manufacturer’s instructions (Glycobuffer 3, 25ºC or 37ºC, overnight). Treated proteins were then immediately incubated with slncRNA and SNA (diluted 1:3) as described in the confocal imaging assay.

### NA treatment of LAMP1

20µg of LAMP1 was incubated with 18µl of NA (The Native Antigen Company, Cat. REC32025-500) or a 9-fold dilution, corresponding to 1:2 and 1:18 enzyme:LAMP1 molar ratios in NA buffer (50mM MES, 2mM CaCl_2_, 150mM NaCl, pH 6.5) at 37ºC overnight. Treated proteins were then immediately incubated with slncRNA and SNA (diluted 1:3) as described in the confocal imaging assay.

### Influenza virus entry assay and analysis

24h pre-infection, 2×10^4^ A549 cells^49^ were seeded in a 96-wells plates in 100µl of DMEM growth media (Sigma, cat. D5796-500ML, supplemented with penicillin-streptomycin solution (IMBH, cat. L0022-100), FBS (IMBH, cat. S1400-500, and MEM NEAA solution (Sartorius, cat. 01-340-1B)) and allowed to adhere and grow overnight for 90% confluency.

On the day of infection, Endo H-(treated or untreated) LAMP1 decoys (102.6pmol slncRNA and 1044pmol protein) were prepared at room temperature 2h prior to the start of the assay. Decoys were then mixed with either 5, 1, 0.5, or 0.1µl of influenza virions (Human IFV A\H1N1\P.R.\8\34 strain), and each condition was performed in duplicates. For no-treatment we used the same viral volumes without any LAMP1 decoys.

A virus-decoy plate was prepared as follows: 21µl of decoys were added to rows C-D (Endo-) or E-F (Endo+). 21µl of infection media, namely growth media without FBS and with type IX-S trypsin (Sigma, cat. T0303-1G), was placed in wells A-B (untreated) and G-H (uninfected cells). Then, the volumes were completed to 100µl using infection media containing one of the tested viral volumes (columns 1-4, respectively to the tested viral volumes). Uninfected control had 0µl of virus. Finally, the media in the cell plate was removed and the content of the virus-decoy plate was transferred to the cell plate, for a 1.5h incubation at 37°C, 5% CO_2_. At the end of the incubation period, the infection media was removed and replaced with fresh growth media. Infection was allowed to take place for 48h.

Following the period of infection, cells were detached mechanically and transferred to a U-bottom 96-well plate (Greiner, cat. 650101) in growth media. Cells were then centrifuged at 1500rpm, 4 °C for 5min. Media was removed without disturbing the cell pellet in the wells, and cells were resuspended and washed in 100µl FACS buffer (PBSx1, supplemented with 1%w/v BSA (Sigma, cat. A9418-50G)), then centrifuged again. The rinse step was repeated once more, before resuspending cells with 100µl CR6261 primary antibody solution (FACS buffer, supplemented with 1mg/µl of mouse-anti-HA antibody, freshly made). Plates were incubated for 1.5h, 4°C. At the end of the incubation period, cells were washed again before staining with 100µl secondary antibody solution (FACS buffer, supplemented with 1:200 AF488-labelled donkey-anti-mouse antibody (ENCO, cat. 715-545-151) and 1:1000 dilution of live/dead stain (Thermo Fisher, cat. 65-0865-14)). Plates were then incubated for 1h, 4°C protected from light. Finally, cells were washed twice more, and resuspended in 200µl of FACS buffer.

Finally, cells were taken for flowcytometry analysis (Fortessa-II). The flow cytometer was calibrated to compensate for the different labels in the system using compensation beads as follows: for FITC and live/stain dye, dedicated beads (Thermo Fisher, cat. 01-111142) were used as follows: 1µl of beads and 1µl of antibody with the same label were added to 1ml of FACS buffer. Beads were then centrifuged and 800µl were removed before resuspension. The same process was used for unlabeled beads. For mCherry compensation, we used 1 drop in 1ml of dedicated beads (Thermo Fisher, cat. A54743) in 1ml FACS buffer.

Analysis of results was performed using custom MATLAB code. Matlab’s fit function was used to estimate the IC_50_ values (Figure 5B+C), using the following model (equation #16) for the dose response *I*:

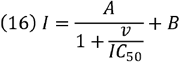

where *A* is the difference between the infected percentages of the highest infection level and the mean of the false positive (FP) signal for uninfected cells (%_high_-%_mean FP uninfected._), *B* is the intercept for the infection, dependent variable (%_mean FP uninfected_), *IC*_50_ is the half-maximal volume needed to get the highest level of infection, *v* is the independent variable (viral volume), *n* is the Hill coefficient, and *I* is the percent of infected cells in a sample. Test article amount was used as µl, and therefore the *IC*_50_ units are the same. The starting point for the fit was *A* =the difference between maximal mean percent between replicates and minimal mean percent of infected cells in the samples; *B* =the maximal mean percent of infected cells; *IC*_50_=the average between the maximal and minimal mean percents of infected cells, *n*=1. Due to the low number of replicates and few test viral volumes, we compiled all 16 possible sets of 4-points using the two replicates per Hill curve. We fitted each set to get its IC_50_ value and report the mean IC_50_ in Figure 5C. The error bars mark the standard deviation of the 16 calculated IC_50_ values. To determine whether there is a significant difference between each 2 conditions, we performed 256 permutations by pooling the four data points per viral volume (two replicates from each curve) and making all possible reassignments of these measurements between the two conditions, refitting the dose–response curves for each reassignment and recalculating the IC50 difference to generate a null distribution. The p-value was then calculated as the proportion of permutations in which the absolute IC_50_ difference was equal to or greater than the observed IC50 difference between the original, unpermuted curves.

### Statistics and reproducibility

The statistical analysis was calculated using a custom Matlab code. For in vitro image analysis experiment, the data for each protein was obtained from several hundred granules. For distribution comparison, a standard two-sided student’s t-test was used.

The infection experiment was carried out in biological duplicates. A Mann-Whitney statistical test was used to assess intra-viral-volume statistical difference and significance.

No statistical method was used to predetermine sample size. No data was excluded from the analysis. The experiments were not randomized. The investigators were not blinded to allocation during experiments and outcome assessment.

## Supporting information

Supplemental Table for Protein Sequences

Movie S1 - Lack of phase transition for bacterial tdPCP

Movie S2 - A phase transition for mammalian tdPCP

Movie S3 - 3D represeentation of a phase transition field of view

## Data availability statement

The data supporting the findings of this study is publicly available at FigShare under accession code 10.6084/m9.figshare.31198237.

## Code availability statement

The image analysis code for the SNA binding assay can be found at https://doi.org/10.5281/zenodo.18421747. The image analysis code for the glycosidase assay can be found at https://doi.org/10.5281/zenodo.18421804. The code for the infection model can be found at https://doi.org/10.5281/zenodo.18421983.

## Author Contributions

**OW** designed and cloned the constructs for LAMP1, GYPA, Fetuin, and the NXST variants, as well as designed and carried out the experiments and analysis for all of the data. **OW** and **RA** conceived the viral infection model. **NG** contributed to part of the computational model, assisted in slncRNA synthesis, and inferred the protein structures using AlphaFold. **OW** and **SG** designed the construct for ACE2. **TS** assisted in the Endo H and NA assays. **SG** designed and cloned tdPCPm, cloned NXST1b, and guided the microscopy experiments and image analysis. **RA** supervised the study. **OW** and **RA** wrote the manuscript.

## Acknowledgements

We thank the Life Sciences and Engineering (LS&E) Infrastructure Center at the Technion (Dr. Nitsan Dahan and Dr. Yael Lupu-Haber from the microscopy core facility) for their excellent technical assistance with confocal microscopy imaging. We thank Tomer Maman for her assistance in the lab preparatory work for this research. We thank Asst. Prof. Yotam Bar-On and Dr. Orly Kladnitsky from the Faculty of Medicine at the Technion for the IFV strain and the cells and materials required for the viral entry assay. We would like to express our gratitude to Amir Grau from the Cytometry Center at the Rappaport Faculty of Medicine Biomedical Core Facility (BCF) for assisting us with the flow cytometry. This work was supported by the European Union’s Horizon 2020 Research and Innovation Programme under grant agreements no. 851615 and no. 851065.

