## Supplemental Figures for "High avidity phase-separated RNA-protein sialogranules sense lectins and inhibit influenza infection"

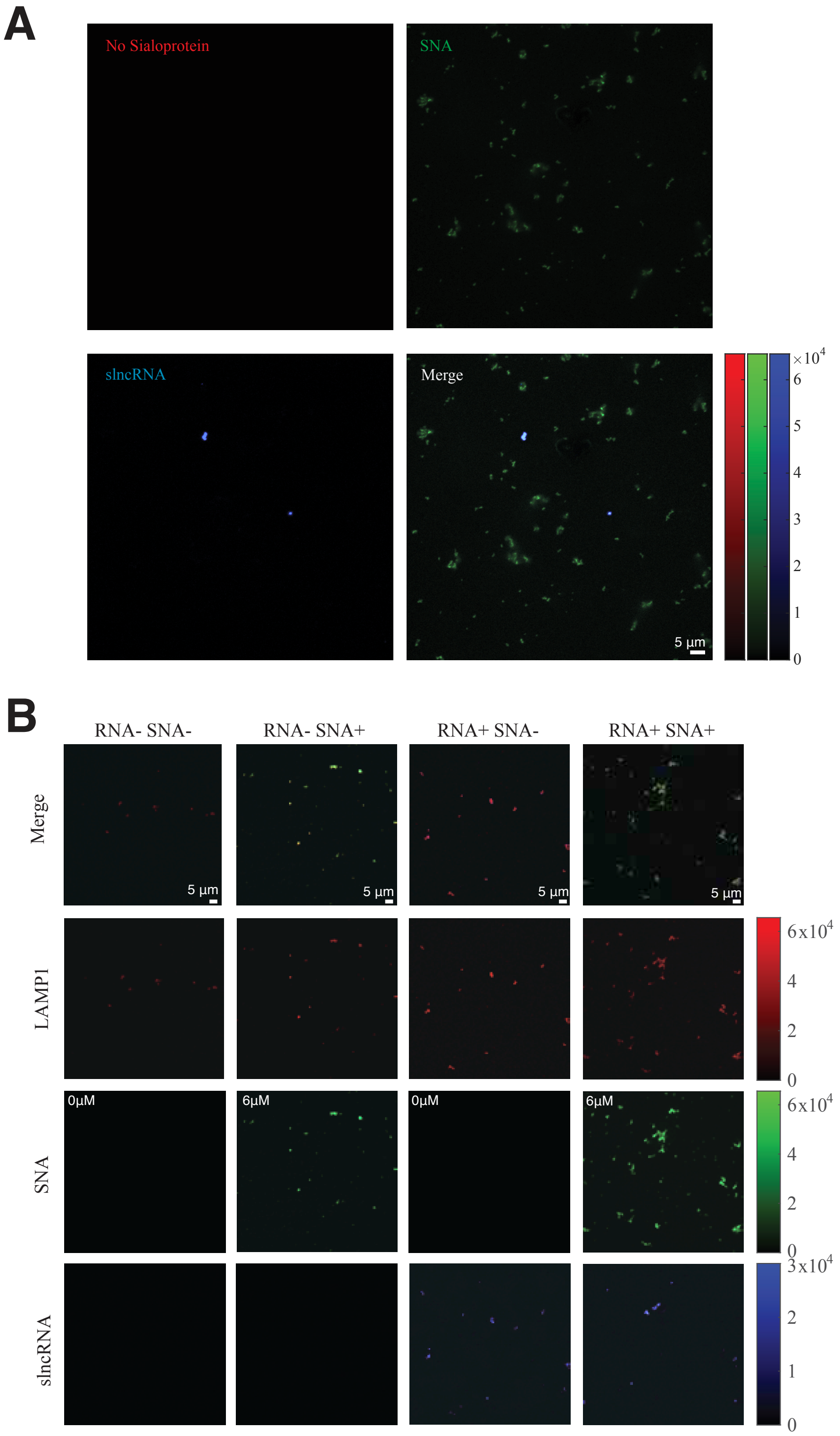
A
No Sialoprotein
SNA
slncRNA
Merge
5 µm
×10^4
6
5
4
3
2
1
0
B
RNA- SNA-
RNA- SNA+
RNA+ SNA-
RNA+ SNA+
Merge
LAMP1
SNA
slncRNA
5 µm
0µM
6µM
6x10^4
4
2
0
3x10^4
2
1
0

**Figure S1 – slncRNA assists in imaging candidate sialoproteins.**

**(A)** slncRNA and SNA cannot interact without the presence of a sialoprotein mediating between them. (Top left) mCherry channel (sialoprotein). (Top right) FITC channel (SNA). (Bottom left) AF405 channel (slncRNA). (Bottom right) Merged channels. Scalebar: 5µm. **(B)** slncRNA presence results in larger biocondensates that form once SNA is added. Additionally, the mCherry signal is strengthened by the presence of the slncRNA, as previously reported[48]. Scalebar: 5µm.

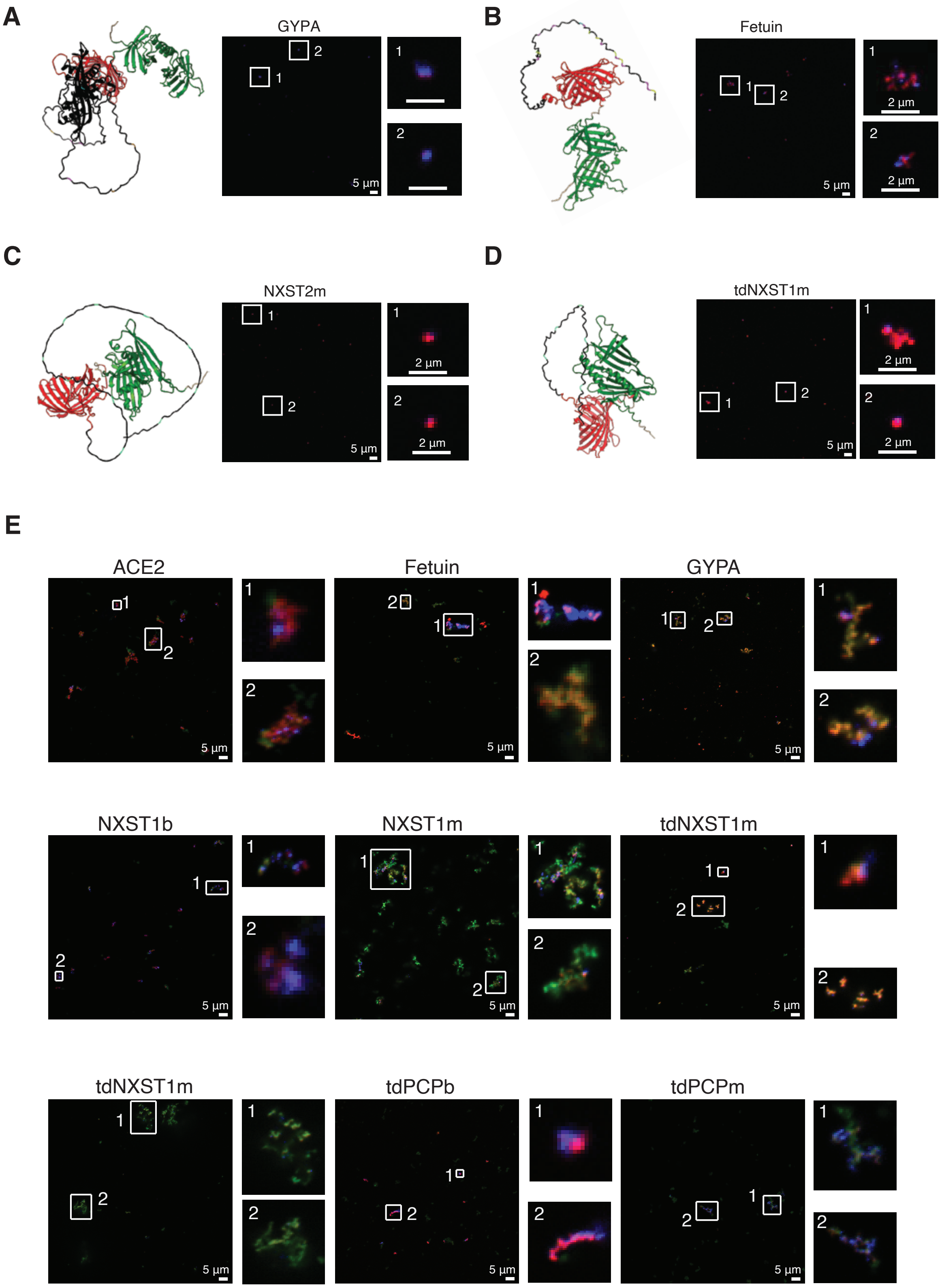

A
GYPA
5 µm
B
Fetuin
2 µm
5 µm
C
NXST2m
2 µm
5 µm
D
tdNXST1m
2 µm
5 µm
E
ACE2
Fetuin
GYPA
NXST1b
NXST1m
tdNXST1m
tdNXST1m
tdPCPb
tdPCPm
5 µm

**Figure S2 – Sialoprotein candidates form granules in the presence of slncRNA and interact with SNA.** Formation of sialogranules for **(A)** GYPA. **(B)** Fetuin. **(C)** NXST2m. **(D)** tdNXST1m. Black, red, and green are sialoprotein, mCherry, and tdPCP domains, respectively. Putative sialylated sites are marked in light blue (asparagine residues, part of the N-X-S/T motif), yellow (serine), or magenta (threonine). Scalebar: 5µm (large FoV) and 2µm (events enlargement). **(E)** Addition of SNA results in the formation of MABs for nearly all sialoprotein candidates. Scalebar: 5µm.

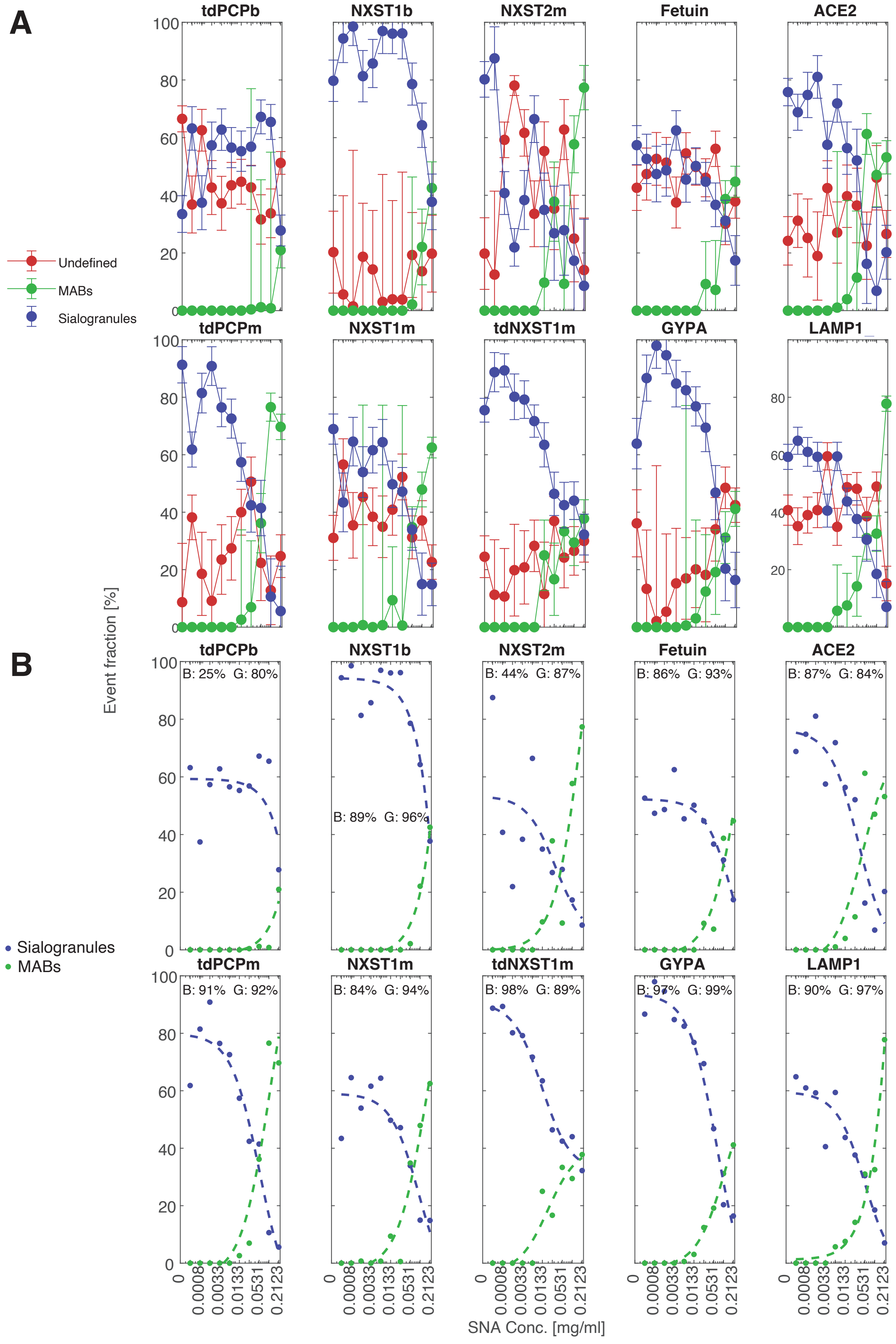
A
B
tdPCPb
NXST1b
NXST2m
Fetuin
ACE2
tdPCPm
NXST1m
tdNXST1m
GYPA
LAMP1
Undefined
MABs
Sialogranules
Event fraction [%]
B: 25% G: 80%
B: 89% G: 96%
B: 44% G: 87%
B: 86% G: 93%
B: 87% G: 84%
B: 91% G: 92%
B: 84% G: 94%
B: 98% G: 89%
B: 97% G: 99%
B: 90% G: 97%
SNA Conc. [mg/ml]

**Figure S3 – Kinetic analysis of SNA binding events. (A)** Mean event frequency (EF) for various sialogranule candidates and MABs (if they are detected). mCherry-positive events are classified as either colocalized with AF405 (RNA, blue), FITC (SNA, green), both (triple label, magenta), or neither (red) (see Methods and Figure 3C). As SNA concentration increases, colocalization with SNA increases while colocalization with RNA decreases. **(B)** Fitted curves of the data in (A).

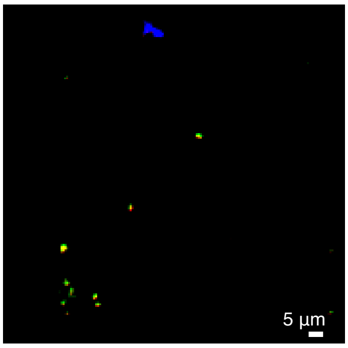


**Figure S4 – Representative field of view of granules made with LAMP1 that was treated with Endo H at 25ºC.** Treatment yielded a recovery in granule event frequency that was lesser than the recovery observed for LAMP1 treated at 37ºC (Figure 4C).

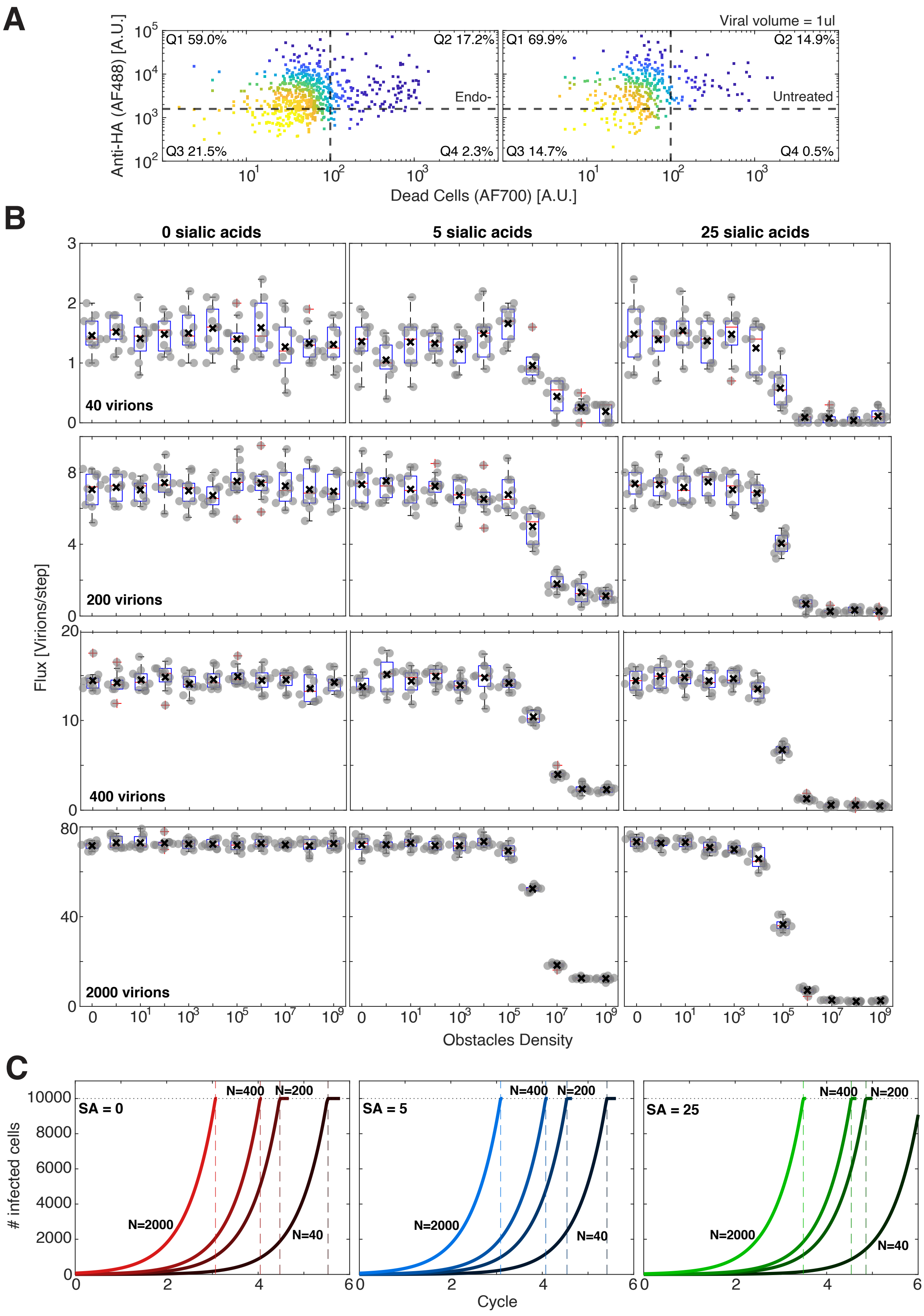
A
Viral volume = 1ul
Q1 59.0%
Q2 17.2%
Q3 21.5%
Q4 2.3%
Endo-
Q1 69.9%
Q2 14.9%
Q3 14.7%
Q4 0.5%
Untreated
Anti-HA (AF488) [A.U.]
Dead Cells (AF700) [A.U.]
B
0 sialic acids
5 sialic acids
25 sialic acids
40 virions
200 virions
400 virions
2000 virions
Flux [Virions/step]
Obstacles Density
C
SA = 0
SA = 5
SA = 25
N=2000
N=400
N=200
N=40
### infected cells
Cycle

**Figure S5 – Raw data from viral entry assay and model simulation. (A)** Representative 2D plots of HA-positive cells (y-axis) and live/dead-stain-negative cells (x-axis) from 1 µl virus mixed with Endo H-untreated LAMP1 decoys (left) and no decoys (right). **(B)** Flux results for 10 runs of the viral infection simulation (see Methods). Mean flux is marked with a black X. A gray dot is a flux result of one simulation. The mean flux results was used to generate the heatmap in Figure 5D, and the distributions of 100k obstacles density was used to generate Figure 5E. **(C)** The predicted number of infected cells stemming from the flux results of each virion count for each of the sialic acid content simulated. Dark-to-light coloring represents increase in viral count. Dashed vertical lines represent the predicted cycles it takes to reach 10k infected cells. Colors correspond to the experimental infection curves in Figure 5B.

**Supplementary Movie S1 – tdPCPb is not affected by the addition of SNA.** Blue – slncRNA. Red – tdPCPb.

**Supplementary Movie S2 – tdPCPm transitions from granular shapes into a liquid-like dense cloud of SNA and sialoproteins, with slncRNA at the periphery.** Blue – slncRNA. Green – SNA. Red – tdPCPm.

**Supplementary Movie S3 – 3D rendering of a LAMP1 granule.** slncRNA (blue) is clustered at the periphery of an SNA (green)-LAMP1 (red) overlap.
